# Binding of Calcium and Magnesium to Cardiac Troponin C

**DOI:** 10.1101/2020.06.14.150854

**Authors:** K Rayani, JT Seffernick, YA Li, JP Davis, AM Spuches, F Van Petegem, RJ Solaro, S Lindert, GF Tibbits

## Abstract

Cardiac troponin C (cTnC) is the Ca^2+^-sensing component of the thin filament. It contains structural sites (III/IV) which bind both Ca^2+^ and Mg^2+^, and a regulatory site (II) that has been thought to bind only Ca^2+^. The latter binding initiates a series of conformational changes that culminate in force production.

We have quantified the interaction between site II and Ca^2+^/Mg^2+^ through Isothermal Titration Calorimetry and Thermodynamic Integration simulations. Direct and competitive binding titrations using wild type and a double mutant that significantly reduces binding to site II demonstrated that physiologically relevant concentrations of both Ca^2+^/Mg^2+^ interact with the same locus. Cytosolic free Mg^2+^ (~1 mM) could occupy a significant population of available site II, as this concentration of Mg^2+^ decreased the affinity for Ca^2+^ 1.4-fold.

Interaction of Mg^2+^ with site II of cTnC likely has important functional consequences for the heart at baseline and in diseased states which decrease or increase availability of Mg^2+^ such as secondary hyperparathyroidism or ischemia, respectively.

## Introduction

Cardiac troponin (cTn) is a heterotrimeric complex that includes components for: Ca^2+^ binding (troponin C – cTnC), inhibition of contraction (troponin I – cTnI), and tropomyosin binding (troponin T – cTnT) (Parmacek and Solaro 2004). Ca^2+^ binding to site II of cTnC is the precursor to a series of structural perturbations in the thin filament (TF) that culminate in a strong force generating reaction between the actin filament and myosin heads (Kawasaki and van Eerd 1972, Murray and Kay 1972, Potter and Gergely 1975, Filatov, Katrukha et al. 1999, Parmacek and Solaro 2004).

cTnC is a 161-amino acid protein composed of 9 helices (N and A-H), which form 4 helix-loop-helix binding motifs (sites I-IV). Within these domains, residues in positions 1 (+x), 3 (+y), 5 (+z), 7 (-y), 9 (-x), and 12 (-z) contain oxygen atoms arranged in a pentagonal bipyramid allowing for coordination of metal cations (**Figure S1**) (Strynadka and James 1989, Yap, Ames et al. 1999, Lewit-Bentley and Rety 2000). Skeletal muscle troponin TnC (sTnC) has 4 functional Ca^2+^ binding motifs (Seamon, Hartshorne et al. 1977, Ebashi, Nonomura et al. 1980). Cardiac TnC (cTnC) has a similar overall structure but a slightly different primary sequence. The insertion of a valine at residue 28, along with the substitutions D29L and D31A have rendered site I of cTnC non-receptive to Ca^2+^ binding (van Eerd and Takahshi 1976, Farah and Reinach 1995).

Ca^2+^ binding to sites III and IV in the C-domain of cTnC occurs with high affinity (~10^7^ M^-1^) (~10x higher than the N-domain) and a slow exchange rate (~100x slower than binding to the N-domain). Given the abundance of contractile filaments throughout cardiomyocytes, cTnC buffers a significant portion of cytosolic Ca^2+^ (Johnson, Charlton et al. 1979, Johnson, Nakkula et al. 1994, Schober, Huke et al. 2012). At resting free cytosolic Ca^2+^ concentrations, sites III and IV are usually saturated with Ca^2+^ (Bers 2000). Mg^2+^ also binds at sites III and IV, but with lower affinity (K_A_~10^4^ M^-1^) (Potter and Gergely 1975). However, the cytosolic concentration of Mg^2+^ allows this cation to compete with and reduce the binding of Ca^2+^ to the “structural” sites (Leavis and Kraft 1978, Robertson, Johnson et al. 1981). The binding of Ca^2+^/Mg^2+^ to sites III and IV alters the structure of TnC and is a prerequisite for thethering to the rest of the TF (Sturtevant 1977, Tikunova and Davis 2004).

The C-domain of cTnC is linked to the N-domain by a linker region composed of a nine turn α-helix (Sundaralingam, Bergstrom et al. 1985, Sia, Li et al. 1997). Within the N-domain (N-cTnC), Ca^2+^ binds the low affinity (~10^-5^ M) site II such that this site is only partially occupied at diastolic free Ca^2+^ concentrations (~0.1 μM) (Cheung, Tillotson et al. 1989). The degree of occupancy is significantly higher at systolic free Ca^2+^ concentrations (~ 0.5 – 1.2 μM) which follow Ca^2+^-induced Ca^2+^-release (Kirschenlohr, Grace et al. 2000). Ca^2+^ binding to site II provides the free energy to allow exposure of a hydrophobic pocket, which is otherwise less favorable (Gifford, Walsh et al. 2007, Bowman and Lindert 2018). Helices B and C (BC domain) move away from helices N, A, and D (NAD domain) to expose the hydrophobic cleft, with the short anti-parallel β-sheet between EF-hands I and II acting as a hinge (Herzberg and James 1985, Slupsky and Sykes 1995, Houdusse, Love et al. 1997). Binding of the “switch peptide” TnI_147-163_ to this pocket facilitates exposure of this hydrophobic region (Sia, Li et al. 1997, Spyracopoulos, Li et al. 1997, Li, Spyracopoulos et al. 1999).

A persistant and under-investigated question is the role of cellular Mg^2+^ in the signaling of activation by cTnC. Of the total cytosolic [Mg^2+^]_i_ (~10 mM), the majority is bound to cellular components such as ATP with only ~0.5 – 1.0 mM being freely available in the cytosol (Romani and Scarpa 1992, Dai, Friedman et al. 1997). In conditions with diminished buffering capacity, such as ATP depleted states, the free [Mg^2+^] can increase significantly (Hongo, Konishi et al. 1994, Tessman and Romani 1998) prior to being extruded from the cell (Laires, Monteiro et al. 2004), but it could also compete with Ca^2+^ for binding to cTnC.

Increase in Mg-ATP in both skeletal and cardiac tissue decreases the Ca^2+^ sensitivity of skinned fibers (Godt 1974, Best, Donaldson et al. 1977, Godt and Morgan 1984). Further evidence has been obtained through fluorescence-based studies of isolated cTnC (Potter, Robertson et al. 1981, Ogawa 1985, Zot and Potter 1987, Morimoto 1991, Francois, Gerday et al. 1993, She, Dong et al. 1998), the Tn complex (Potter, Robertson et al. 1981, Zot and Potter 1987), and reconstituted fibers (Zot and Potter 1987, Allen, Yates et al. 1992) where Mg^2+^ appears to decrease Ca^2+^ sensitivity. In isolated TnC, the K_A_ of the low affinity sites (III/IV) for Ca^2+^ and Mg^2+^ was measured to be on the order of 10^6^ M^-1^ and 10^2^ M^-1^, respectively (Ogawa 1985).

Interaction of Ca^2+^/Mg^2+^ with sites III/IV results in a large change in enthalpy (ΔH), in contrast to changes resulting from site I/II binding which are small by comparison. Detection of heat changes associated with the interactions of metal ions and proteins is both challenging and highly technique dependent such that small changes may be deemed negligible (Yamada 1978, Yamada and Kometani 1982, Kometani and Yamada 1983). Experiments used to study this system decades ago were limited by the technology of the time. In contrast, modern Isothermal titration calorimetry (ITC) is a sensitive method to that can be used to define the thermodynamic parameters of binding without the use of labelling methods that could interfer. Modern ITC allows for the study of single binding sites within isolated protein domains and can be used to detect heat changes as small as 0.1 μcal (Yamada 2003, Wilcox 2008, Grossoehme, Spuches et al. 2010, Sacco, Skowronsky et al. 2012).

We have used ITC to explore the binding of Mg^2+^ and Ca^2+^ to site II at the level of N-cTnC and full-length cTnC. Competitive binding to the N-domain and mutations in the site II caused a reduction in apparent affinity, indicating interaction of both cations with the same locus in the protein. In full-length cTnC, Mg^2+^ competed with and reduced Ca^2+^ binding to all three sites. These findings further corroborate and expand upon what has been shown by a few labs, but the findings are largely ignored by most; the role of Mg^2+^ in modulating the Ca^2+^-sensitivity of force production in cardiomyocytes is one which merits further discussion.

## Methods

### Construct preparation and protein expression

The *TNNC1* gene (Uniprot ID P63316) had previously been cloned into pET21a(+) vector and had a stop codon inserted at residue 90 to create the N-cTnC construct using the Phusion site-directed mutagenesis protocol (Thermo Scientific). This construct was transformed into the BL21(DE3) expression strain. The D76A/D73A construct was made using site-directed mutagenesis carried out by GenScript (New Jersey, USA). Expression and purification of all constructs were carried out as described previously (Stevens, Rayani et al. 2016, Stevens, Rayani et al. 2017). In brief, 100 mL of lysogeny broth (LB) was supplemented with 50 μg/mL ampicillin and a glycerol stock stab and grown over-night at a shaking speed of 250 rpm and 37°C. In the morning, the same conditions were provided to 1 L cultures that were grown for ~3 hrs to an OD_600_ of 0.8 – 1.0 followed by induction with β-D-1-thiogalactopyranoside (IPTG). After 3 hrs, the cells were harvested by centrifugation and stored at −80 °C.

### Protein purification

The cell pellet was thawed and suspended in 50 mM Tris-Cl pH 8.0, 5 mM ethylenediamine-tetraacetic acid (EDTA) and sonicated on ice, 5 times 30 sec each, with intervals of rest between to avoid overheating. The lysate was centrifuged for 15 mins at 30,000 ×g and the supernatant separated twice to remove all cell debris. The supernatant was applied to a fast-flow Q-Sepharose column pre-equilibrated with the suspension buffer and 1 mM dithiothreitol (DTT). The protein was eluted from the column by applying a 180 mL ramp gradient to the same buffer supplemented with 0.5 M NaCl. The gradient was applied using an AKTA FPLC machine that was also used to fractionate the eluted samples. Following analysis by SDS-PAGE, the samples containing N-cTnC were pooled and concentrated using Amicon centrifugal concentrators with a 3 KDa molecular weight cut-off (Millipore).

The full-length protein was purified in the same way, with the addition of a 30% ammonium sulphate following the sonication step. Centrifugation was then used to remove insoluble components and the supernatant was then dialyzed overnight against 4 L of column-buffer containing 50 mM Tris-Cl pH 8.0 and 100 mM NaCl.

The concentrated protein was then applied to a HiPrep 26/60 Sephacryl S-100 column (DEAE FF) (GE healthcare) which was equilibrated with the re-suspension buffer supplemented with 100 mM NaCl. SDS-PAGE analysis of the fractions was used to identify and pool those containing cTnC. The protein was stored at −80 °C prior to pre-ITC dialysis.

### Dialysis and ITC experiments

To generate the apo-state protein, troponin C was first dialyzed against 2 L of 50 mM HEPES pH 7.2, 150 mM KCl, 2 mM EDTA, and 15 mM β-mercaptoethanol (BME), followed by another dialysis against the same buffer. Each of these dialysis steps was completed at 4°C for a minimum of 4 hrs. A third dialysis was performed for a minimum of 16 hrs overnight against 2 L of 50 mM HEPES pH 7.2, 150 mM KCl, and 2 mM EDTA. An extinction coefficient of 1490 M^-1^cm^-1^ and 4595 M^-1^cm^-1^, and a molecular weight of 10.1 kDa and 18.4 KDa were used to determine protein concentration for the N-cTnC and full length cTnC constructs, respectively, by 280 nm UV-vis spectroscopy using a NanoDrop 2000 spectrophotometer (Thermo Scientific). The final dialysis buffer was used to dilute the protein samples to a final concentration of 200 μM for the N-terminal construct and 150 μM for full length cTnC as described previously (Stevens, Rayani et al. 2017).

### Experimental Protocols

#### Full-length cTnC

Standard 1.0 M CaCl_2_ and MgCl_2_ stock solutions (Sigma, USA) were serially diluted in the final dialysis buffer to produce 6 mM Ca^2+^ and 40 mM Mg^2+^ for the full length cTnC titrations. 6 mM Ca^2+^ was titrated into 100 μM apo-state full-length human cTnC as the baseline condition. The data were fit with a two sets-of-binding-sites model. The same amount of protein was diluted in the ITC buffer and used for all subsequent conditions that were fit with the same model. Supra-physiological concentrations Ca^2+^ were selected for the pre-incubation experiments and justified as the goals of these experiments were not violated. The goals of these experiements were to explore possible competition between Ca^2+^ and Mg^2+^ in binding to cTnC and N-cTnC. Higher concentrations could thus be used to observe clearer results. Also, the amounts of cTnC and N-cTnC are above what is available in the cell and the affinity of their binding sites is lowered as they exist in isolation in these experiments ie. their affinities would be elevated by the presence of other cTn complex proteins. The concentrations of Mg^2+^ pre-incubated (1 and 3 mM) is not far beyond what would be expected in normal cellular conditions.

#### N-terminal cTnC

The same standards were used to produce 4 mM Ca^2+^ and 20 mM Mg^2+^ titrants for the N-cTnC experiments. 4 mM Ca^2+^ was titrated into 200 μM apo-state N-cTnC as the baseline condition with subsequent titrations using the same amount of protein. The isotherms were all fit with a single binding site model.

#### Titrations

The ITC experiments were carried out in a MicroCal ITC200 instrument (Malvern, UK). Repeat titrations were used to ensure reproducibility. The sample cell was set at 25 °C, 200 μL of the protein was loaded and the experiment was carried out at the same temperature. For the N-terminal constructs, 19 injections of the titrant were used with the first being a dummy injection of 0.8 μL and the subsequent 18 injections, 2 μL each. For the full-length constructs, the same volume of sample was titrated with a dummy injection of 0.5 μL and 38 injections of 1 μL. The time interval between injections was 120 sec and stirring speed was set at 1,000 rpm throughout each experiment.

### Analysis of results

Titration data obtained from ITC were fit using Origin 8.0 (OriginLab, Northampton, MA) to calculate the thermodynamic parameters using a least-squares algorithm by the software. In this method, if multiple ligands were simultaneously present in the reaction mixture, an “apparent affinity” was determined for the injected titrant. Origin also allows for fitting of more complex models of interactions which were utilized in the case of multiple binding sites for the full length cTnC experiments.

When fitting the data for the N-cTnC constructs (apart from the Ca^2+^ into Apo-state N-cTnC condition), the N associated with each interaction was necessarily constrained to equal 1.00 to facilitate curve fitting without altering protein concentration. The baseline condition was repeated daily to monitor fluctuations in concentration of properly folded and functional protein.

Protein concentration plays a large role in determination of affinity. The concentration of the titrant may be affected by pipetting errors, albeit this effect is normally minimal. The ratio of the ligand to titrant in the single binding site condition (as given by the stoichiometry – n) is a measure of the functional moles of protein and was approximately 1.00 in all the N-cTnC titrations. Given the method of concentration determination, the number of binding sites, cooperativity, and the variable binding strength of each titrants, the N cannot be used in the same way for the full-length cTnC experiments. Therefore, the values presented can be compared between conditions, but care should be taken when comparing these to other systems. Ease of manipulation of the N-cTnC/cTnC system contrasts with those that include the cTn/TF. Thus, the binding parameters measured here may not translate in absolute term when cTnC is incorporated into a more complex system.

### Thermodynamic Integration (TI)

Starting from the representative model of PDB:1AP4 (Spyracopoulos, Li et al. 1997), that contains N-cTnC with a single Ca^2+^ ion bound, the system was solvated with a 12 Å padded TIP3P water box and neutralized with Na^+^ in Amber16 (D.A. Case, R.M. Betz et al. 2016). The system was also prepared similarly for only the Ca^2+^ ion in a 12 Å padded TIP3P water box. The alchemical thermodynamic cycle used for ligand binding was described in detail previously (Leelananda and Lindert 2016). In short, TI was performed using the following three steps for Ca^2+^ in protein: turn on restraints, turn off charge, and turn off van der Waals forces. The specific distance restraints used in all systems can be found in **Table S1**. Additionally, TI was performed for the following two steps for Ca^2+^ in water: turn off charge and turn off van der Waals forces. Each step of the thermodynamic cycle was performed with the coupling parameter (λ) ranging from 0.0-1.0 in increments of 0.1. For each simulation, the system was minimized (2000 cycles) and heated (0.5 ns) before the 5 ns production run at 300 K using the ff14SB force field (Maier, Martinez et al. 2015). These calculations were also performed on the D67A/D73A mutated system. The mutations were imposed on the 1AP4 representative model using PyMOL (L DeLano 2002).

For the calculation of Mg^2+^ binding affinity, Ca^2+^ was replaced with Mg^2+^ in the 1AP4 representative model since no Mg^2+^-bound N-cTnC structure was available in the protein databank. In order to generate more accurate restraints and starting coordinates for the TI calculations, a minimization was performed on the structure in Amber (2000 cycles). Following the minimization, TI simulations were run similarly as for Ca^2+^. However, due to previously documented errors in the default Mg^2+^ parameters, the ΔG_solvation_-optimized Mg^2+^ parameters from Li et. al. were used (Li, Roberts et al. 2013, Panteva, Giambasu et al. 2015). These calculations were also performed on the D67A/D73A mutated system.

To calculate absolute binding affinities for the ions, the change in free energy (ΔG) was calculated for each step in the thermodynamic cycle by integrating the potential energy with respect to the coupling parameter, λ (Shirts, Mobley, et al. 2010). Two corrections were made to these calculated ΔG values. The first correction was necessary due to the introduction of the distance restraints (as described in Boresch et al.) which quantified the free energy cost of restraining the ion to the binding site (Boresch, Tettinger et al. 2003). The second correction was performed to correct the charged system (as described in Rocklin et al.) to revise the free energy for the fact that the system is charged during the disappearance of the charged ions (Rocklin, Mobley et al. 2013). The overall ΔG of binding was the change in free energy between the ions in complex with the protein (ion in protein steps 1, 2, and 3) and the ions in water (ion in water steps 1 and 2). For each system, 5 independent runs were performed, and results were averaged.

## Results

### N-terminal cTnC

### Ca^2+^ and Mg^2+^ binding to apo-state N-cTnC

The interaction of N-cTnC with either Ca^2+^ or Mg^2+^ was found to be associated with a positive ΔH, so the interaction is driven by entropy (**Figure 1**) consistent with previously published data (Skowronsky, Schroeter et al. 2013, Tanaka, Takahashi et al. 2013, Stevens, Rayani et al. 2016, Stevens, Rayani et al. 2017). This supports our interpretation that the endothermic component between full-length cTnC and Ca^2+^ is due to binding to the N-doman.

**Figure 1 -.**
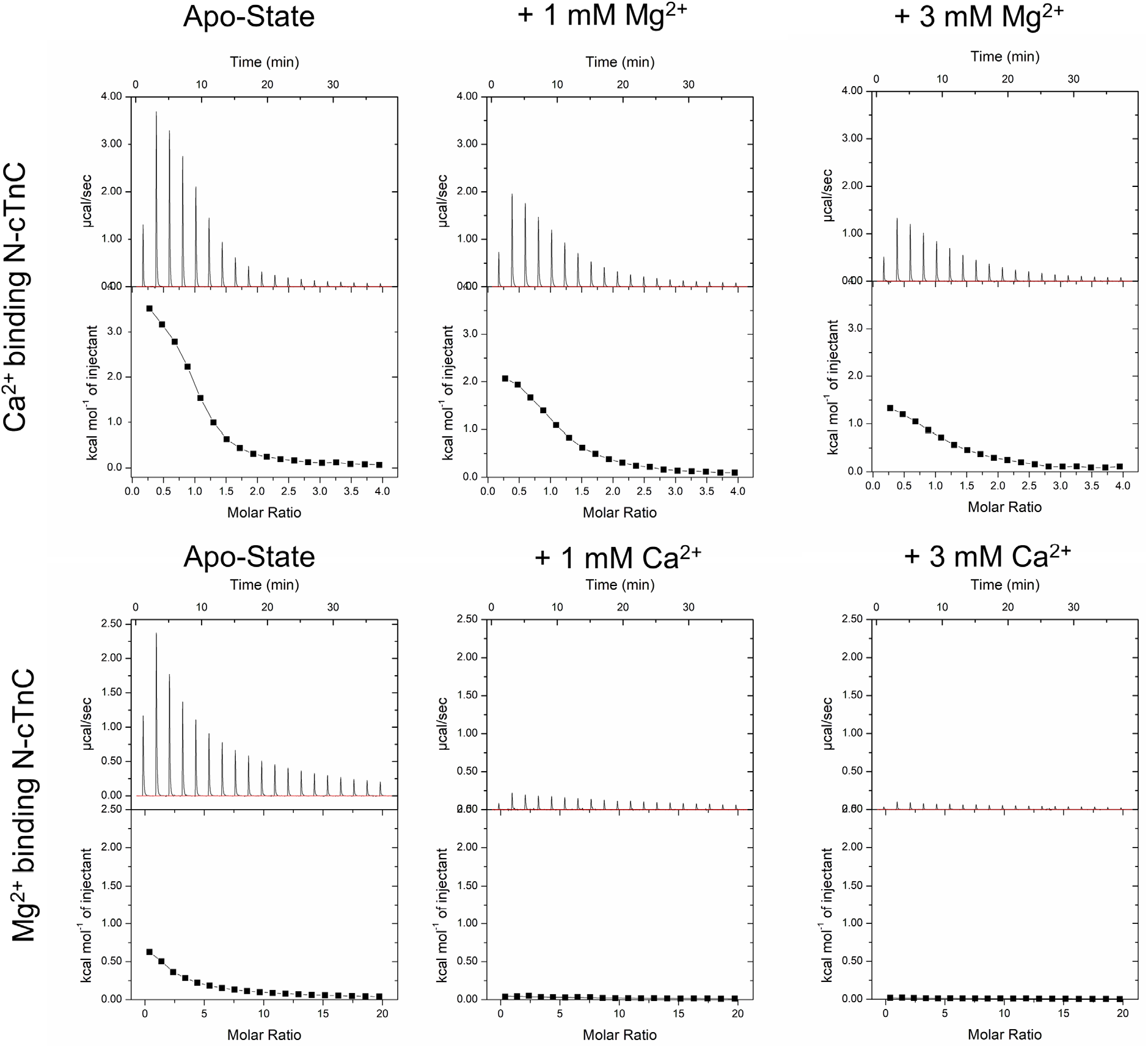
Representative isotherms of Ca^2+^ and Mg^2+^ binding to to apo-state and pre-incubated N-cTnC. Upper panels show the heat recorded during the titration and lower panels plot the integrated heat signal against the molar ratio of titrant added. The top row shows the titration of 4 mM Ca^2+^ into apo-state N-cTnC, followed by the same titration into 1 mM and 3 mM Mg^2+^ pre-incubated N-cTnC. The bottom row shows the titration of 20 Mg^2+^ into apo-state N-cTnC, followed by 1 and 3 mM Ca^2+^ pre-incubated N-cTnC. The fitting of integrated heats was achieved using a single binding site model.

The affinity of N-cTnC for Ca^2+^ (K_d_ = 15.2 ± 0.5 μM) was found to be more than 42.9-fold greater than for Mg^2+^ (K_d_ = 652.8 ± 28.4 μM) difference (**Figures 1 and 2; Table S2**).

**Figure 2 -.**
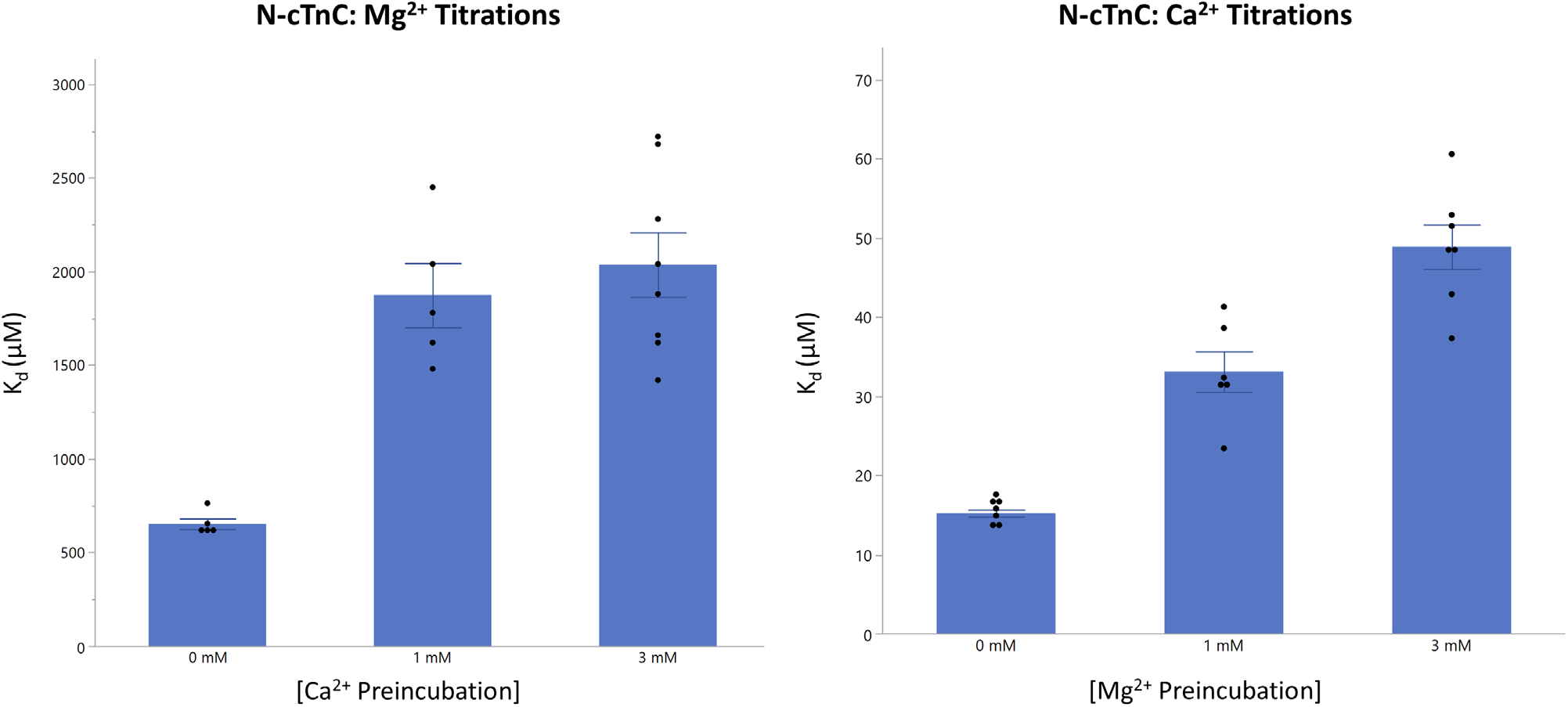
Binding of Ca^2+^ and Mg^2+^ to apo-state and pre-incubated N-cTnC. Left panel: The affinity of site II for Mg^2+^ is compared in the apo-state and with Ca^2+^ preincubation in N-cTnC; Right Panel: The affinity of site II for Ca^2+^ is compared in the apo-state and with Mg^2+^ preincubation in N-cTnC. Statistical differences were assessed through ANOVA followed by Tukey’s post hoc test. Ca^2+^ titrations were not significantly different but the titration of Mg^2+^ into N-cTnC pre-incubated with 1 mM/3 mM Ca^2+^ was signiferently different from the apo-state titration.

The ΔH of the Ca^2+^-N-cTnC interaction was significantly greater (3.82 ± 0.04 kcal*mol^-1^) than that with Mg^2+^ (2.64 ± 0.10 kcal*mol^-1^). Moreover, the entropic contribution for the Ca^2+^ titrations (T*ΔS = 10.39 ± 0.03 kcal*mol^-1^) was more favorable than the Mg^2+^ titrations (T*ΔS = 6.99 ± 0.07 kcal*mol^-1^) (**Table S2**).

As expected, the affinity of Ca^2+^ binding to apo-state N-cTnC was also found to be characterized by the lowest observed dissociation constant (K_d_) (15.2 ± 0.5 μM) of all the titration conditions carried out, including pre-incubation (**Figure 2 and Table S2)**.

### Mg^2+^ binding to Ca^2+^ pre-incubated N-cTnC

To investigate Mg^2+^ binding, apo-state N-cTnC was pre-incubated with increasing concentrations of Ca^2+^ (0, 1, and 3 mM), then titrated with 20 mM Mg^2+^ (**Figure 1 and 2; Table S2**). Mg^2+^ binding the apo-state N-cTnC occurred with significantly lower affinity (652.8 ± 28.4 μM) than the Ca^2+^ titration (15.2 ± 0.5 μM). Moreover, the change in enthalpy in these conditions was significantly lower as increasing amounts of Ca^2+^ was pre-incubated. Titration of Mg^2+^ into apo-state protein yielded a ΔH = 2.64 ± 0.10 kcal*mol^-1^, an order of magnitude lower than Ca^2+^ into apo-protein which liberated 3.82 ± 0.04 kcal*mol^-1^. Moreover, the K_d_ values were 1870.0 ± 171.5 μM and 2037.5 μM ± 172.2 μM for the 1 mM and 3 mM Ca^2+^ conditions, showing a decrease in affinity with increasing concentrations of Ca^2+^ pre-incubated with the protein sample and a more than 2 orders of magnitude lower affinity compare to the Ca^2+^ into WT condition. The significant reduction in affinity, ΔH, and increasingly smaller ΔS associated with higher Ca^2+^ pre-incubation suggests that both metal cations may be binding to the same EF-hand binding motif in site II of N-cTnC.

### Ca^2+^ binding to Mg^2+^ pre-incubated N-cTnC

Apo-protein pre-incubated with Mg^2+^ was titrated with Ca^2+^ to assess the “apparent” affinity of the protein for Ca^2+^ when the site might be occupied with the other divalent cations. As expected, increasing the Mg^2+^ concentration reduced the amount ΔH associated with binding from 3.82 ± 0.04 kcal*mol^-1^ in the apo titration to 1.73 ± 0.05 kcal*mol^-1^ in the 3 mM Mg^2+^ pre-incubated construct. The binding affinity similarily changed from 15.2 ± 0.5 μM to 48.9 ± 2.8 μM. The Ca^2+^ affinity was lower compared to the apo-N-cTnC binding condition and decreased with higher concentrations of Mg^2+^ (**Figure 2 and Table S2**).

### Ca^2+^ and Mg^2+^ binding to apo-D67A/D73A N-cTnC

Point mutations (D67A and D73A) were made (**Figure S2**), affecting two known Ca^2+^ coordinating residues in site II. Binding of both divalent cations was reduced by these mutations but the K_d_ was still lower for Ca^2+^ binding (180.3 ± 16.2 μM) compared to Mg^2+^ binding (1148.6 ± 95.0 μM) (**Figures 3 and 4; Table S2**). Compared to the WT, Ca^2+^ binding was reduced 11.9-fold and Mg^2+^ binding was reduced 1.8-fold by the double mutation. The change in affinity was significantly higher for the Ca^2+^ titrations but statistically indifferent for Mg^2+^.

**Figure 3 -.**
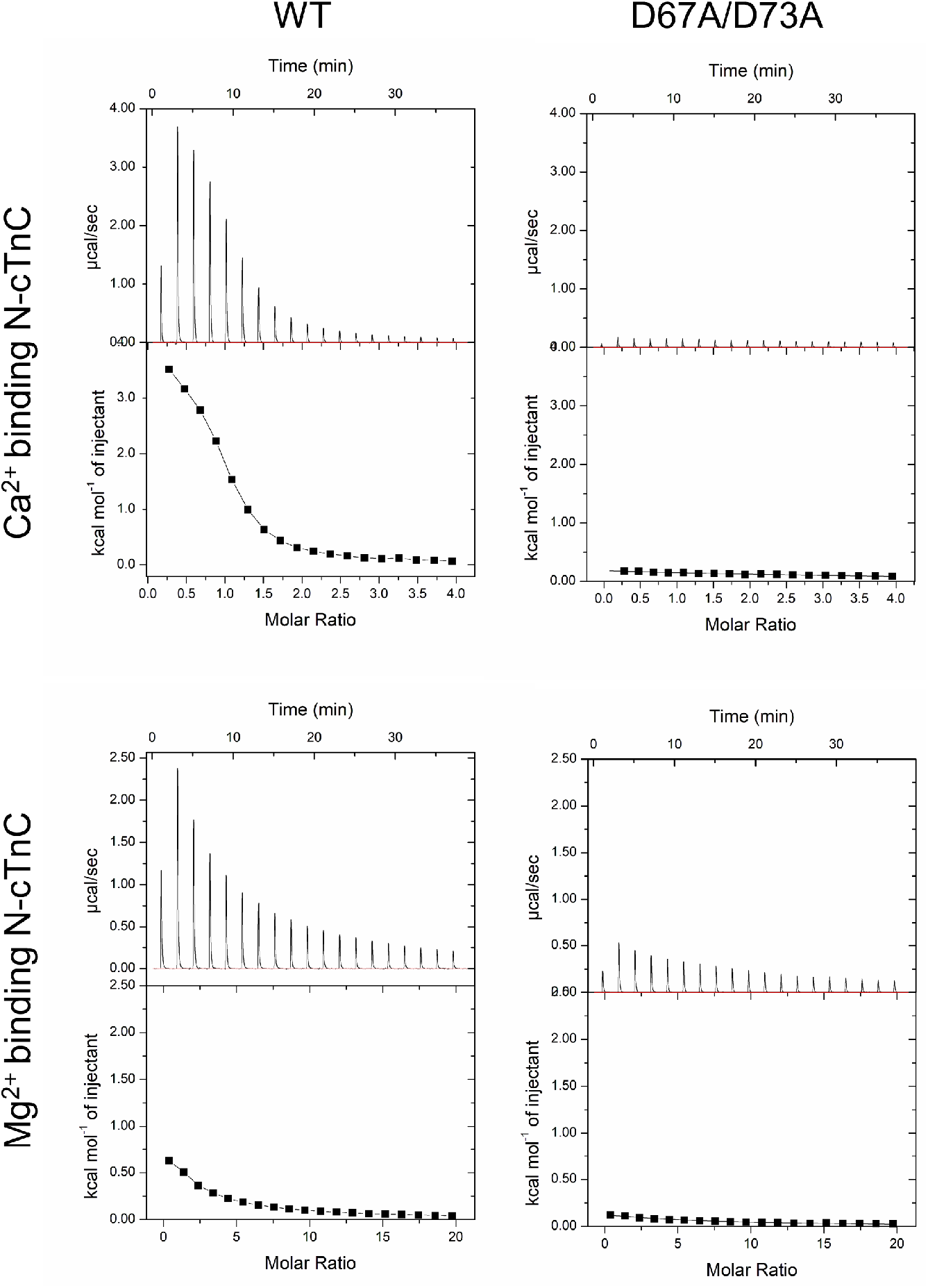
Representative binding isotherms for binding of Ca^2+^ and Mg^2+^ WT and D67A/D73A N-cTnC. The top row shows the titration of 4 mM Ca^2+^ into apo-state N-cTnC, followed by the same titration into the D67A/D73A mutant. The bottom row shows the titration of 20 mM Mg^2+^ into apo-state N-cTnC, followed by the same titration into the D67A/D73A mutant. The fitting of integrated heats was achieved using a single binding site model.

**Figure 4 -.**
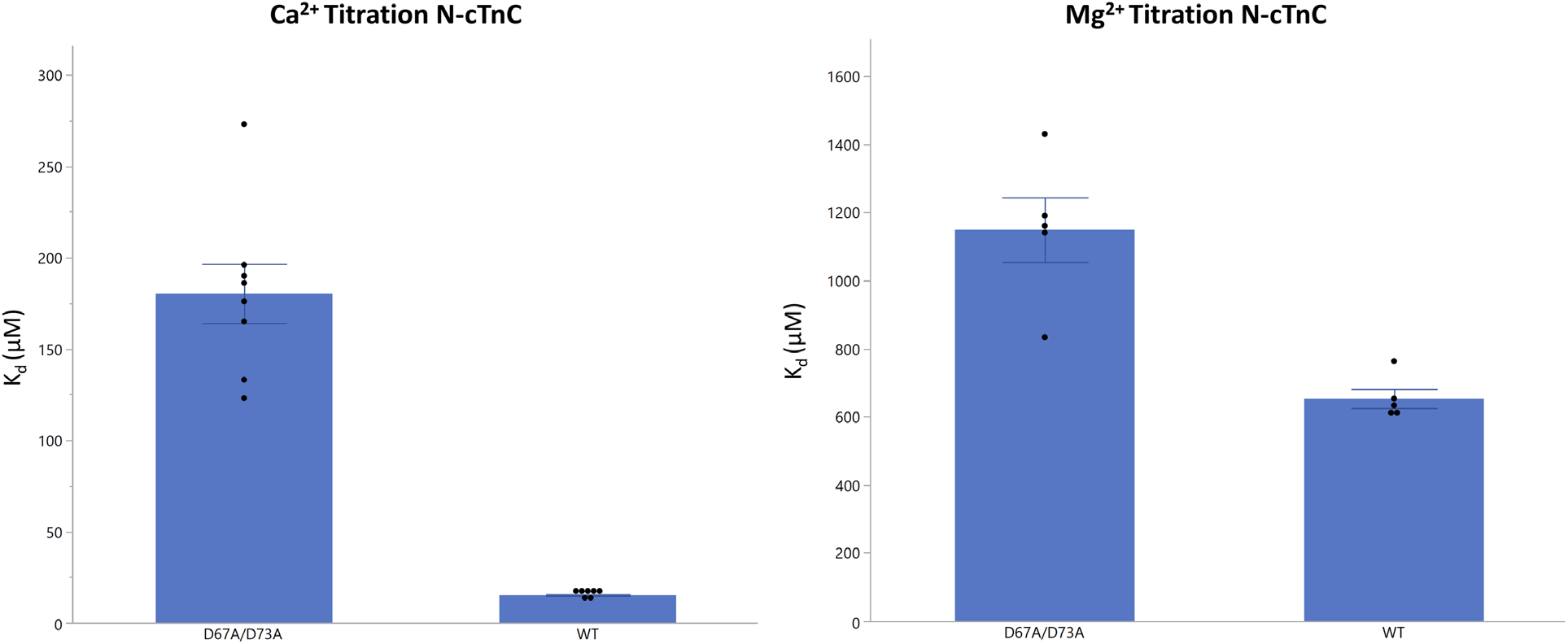
Binding of Ca^2+^ and Mg^2+^ to WT and D67A/D73A N-cTnC. The effect of the D67A/D73A on Ca^2+^ and Mg^2+^ binding is assessed. The affinity of binding for both cations to N-cTnC was lower when comparing the mutant and the WT. The effect on Ca^2+^ binding was more pronounced (11.9-fold reduction) compared to Mg^2+^ (1.8-fold reduction) but this is reconcilable with the number of coordinating residues needed to bind Ca^2+^ (6) vs. Mg^2+^ (5); having 4 available coordinating residues was expected to affect Ca^2+^ binding to a greater extent.

### Ca^2+^ and Mg^2+^ binding affinities from Thermodynamic Integration

Thermodynamic Integration (TI) was performed to calculate absolute binding affinities for the ions in the following systems: Ca^2+^ to WT N-cTnC, Ca^2+^ to D67A/D73A N-cTnC, Mg^2+^ to WT N-cTnC, and Mg^2+^ to D67A/D73A N-cTnC. The average calculated binding affinities over 5 independent runs were −6.9 ± 1.3, −4.5 ± 2.4, −0.6 ± 2.8, and +0.4 ± 2.3 kcal*mol^-1^, respectively. The TI-determined Ca^2+^ binding affinities were in good agreement with the ITC data. While the calculated absolute Mg^2+^ binding affinities were not in perfect agreement with the ITC data, they did show that Mg^2+^ had a weaker binding affinity than Ca^2+^ for all systems (−6.57 to −4.38 kcal*mol^-1^ and −6.9 to −0.6 kcal*mol^-1^ for ITC and TI, respectively for WT system and −5.12 to −4.02 kcal*mol^-1^ and −4.5 to +0.4 kcal*mol^-1^ for ITC and TI, respectively for D67A/D73A system). Additionally, between the Mg^2+^ binding affinities, the binding affinity was consistently weaker for the D67A/D73A mutation. The ΔΔG values comparing ΔG between WT and D67A/D73A systems were similar for ITC and TI (0.36 kcal*mol^-1^ and 1.0 kcal*mol^-1^, respectively).

**Table 1 -.** Average calculated binding affinities for each Thermodynamic Integration system

| System | $\Delta G_{TI}$ (kcal*$\text{mol}^{-1}$) |
| --- | --- |
| $\text{Ca}^{2+}$ to WT | $-6.9 \pm 1.3$ |
| $\text{Ca}^{2+}$ to D67A/D73A | $-4.5 \pm 2.4$ |
| $\text{Mg}^{2+}$ to WT | $-0.6 \pm 2.8$ |
| $\text{Mg}^{2+}$ to D67A/D73A | $+0.4 \pm 2.0$ |

Averages were calculated over 5 independent runs

### Full length cTnC

### Ca^2+^ binding to apo-state full length cTnC

Binding of Ca^2+^/Mg^2+^ to site II is characterized by an endothermic interaction as indicated by our titrations on the N-terminal domain in this and previous publications (Stevens, Rayani et al. 2016, Stevens, Rayani et al. 2017). From this, and ITC work on full-length cTnC by others (Johnson, Fulcher et al. 2019), we can deduce that the exothermic interactions seen above (**Figure 5**) result from interactions with site III/IV. The data (**Figure 5 and Table S3**) show that Mg^2+^ binds to apo-state full-length cTnC at two distinct sets of sites.

**Figure 5 -.**
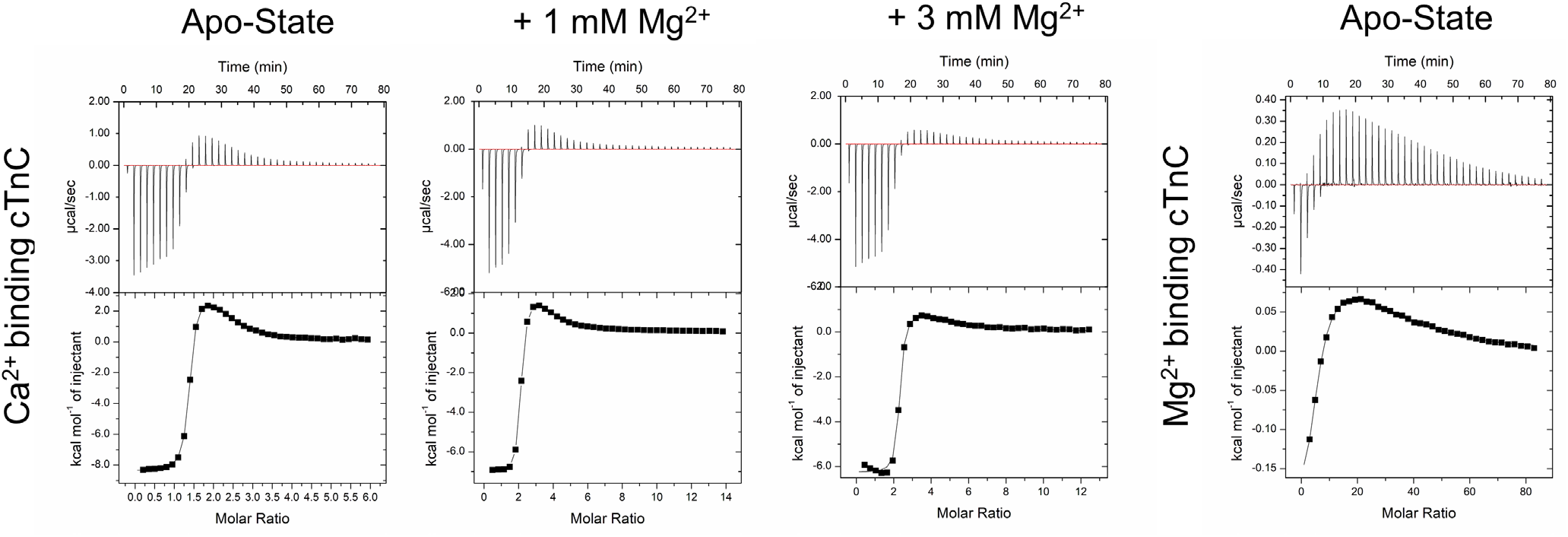
Representative isotherms for binding of Ca^2+^ and Mg^2+^ to full length cTnC. Representative isotherms for each full-length cTnC titration condition are shown. Upper panels show the heat recorded during the titration and lower panels plot the integrated heat signal against the molar ratio of titrant added. A two sets of binding sites model was used to fit the integrated heats. From left to right, the panels show: the titration of 6 mM Ca^2+^ into: apo-state full length cTnC, 1 mM Mg^2+^ incubated cTnC, and 3 mM Mg^2+^ incubated cTnC. The isotherms can be used to visually determine the decreased amount of binding to site II in the presence of increasing Mg^2+^. In the right most panel of this figure, the binding of Mg^2+^ to apo-state full-length cTnC occurs at two sets of different sites as seen in the isotherm which contains both exothermic and endothermic components.

The binding of Ca^2+^ to sites III/IV occurred with an apparent K_d_ of 0.12 ± 0.02 μM, characterized by an exothermic component (ΔH = −8.12 ± 0.07 kcal*mol^-1^) with a positive change in entropy (T*ΔS = 1.24± 0.07 kcal*mol^-1^). In the same full-length construct, the K_d_ associated with binding of Ca^2+^ to site II was 22.7 ± 0.5 μM significantly lower binding affinity. It also had positive ΔH (3.71 ± 0.06 kcal*mol^-1^) and was entropically driven (T*ΔS = 10.0 ± 0.07 kcal*mol^-1^) (**Figure 5 and 6; Table S3**).

**Figure 6 -.**
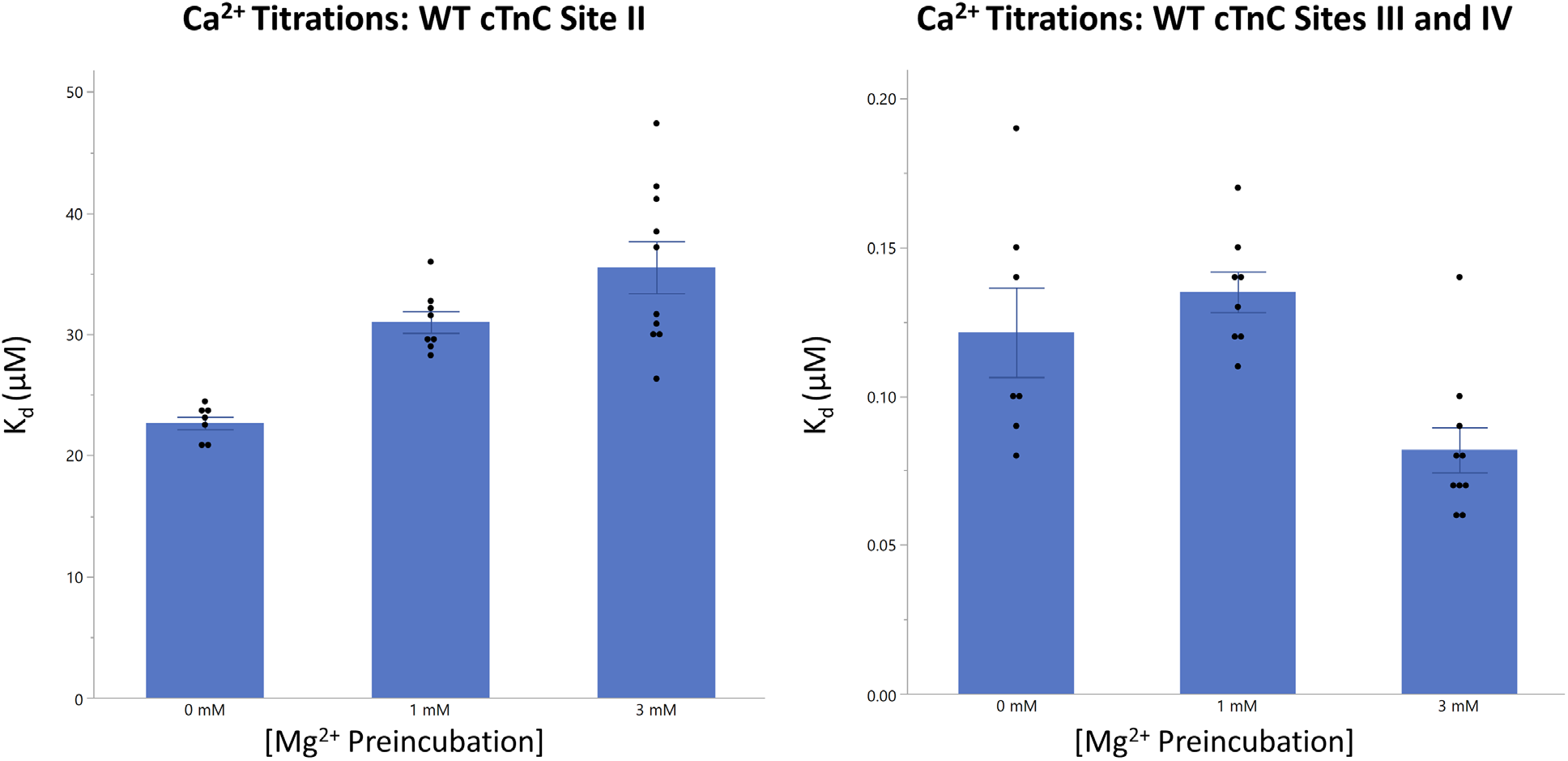
Binding of Ca^2+^ and Mg^2+^ to apo-state and pre-incubated full length cTnC. Left panel: The affinity of site II for Ca^2+^ is compared in the apo-state and with Mg^2+^ preincubation in full length cTnC; Right Panel: The affinity of sites III/IV for Ca^2+^ is compared in the apo-state and with Mg^2+^ preincubation in full length cTnC. At site II, preincubation with 1 and 3 mM Mg^2+^ caused a significant reduction in the affinity for Ca^2+^ binding. At sites III/IV, preincubation with 1 mM Mg^2+^ decreased affinity for Ca^2+^ binding (not statistically significant), while 3 mM Mg^2+^ unexpectedly increased the affinity for Ca^2+^ binding at these sites. Statistical differences were assessed through two-way ANOVA followed by Tukey’s post hoc test.

### Mg^2+^ binding to apo-state full length cTnC

Mg^2+^ binding to site II (K_d_ = 406.1 ± 7.9 μM) and sites III/IV (K_d_ = 16.7 ± 0.7 μM) was characterized by a positive ΔH (0.091 ± 0.001 kcal*mol^-1^) and negative ΔH (−0.23 ± 0.01 kcal*mol^-1^), respectively (**Figure 5 and Table S3**). Based on these enthalpy values, significantly greater amounts of Mg^2+^ binding occur at the C-terminus in comparison to the N-terminus.

The interaction of Mg^2+^ with sites III/IV is two orders of magnitude weaker than that seen for Ca^2+^. The interaction with site II and sites III/IV were both entropically favorable (T*ΔS = 4.71 ± 0.01 kcal*mol^-1^ and T*ΔS = 6.28 ± 0.03 kcal*mol^-1^) and resulted in spontaneous interactions (ΔG = −4.62 ± 0.11 kcal*mol^-1^ and ΔG = −6.51 ± 0.31 kcal*mol^-1^). These all differed significantly from those observed for Ca^2+^ binding, p<0.05.

### Ca^2+^ binding to Mg^2+^ pre-incubated full length cTnC

At greater concentrations, Mg^2+^ occupied a greater proportion of binding sites and limited binding Ca^2+^ to cTnC at all sites (**Figure 5 and 6; Table S3**). Binding of Ca^2+^ to site II was significantly reduced by pre-incubation with 1 mM and 3 mM Mg^2+^ as indicated by an increase in K_d_ and a lowering of the ΔH. Binding of Ca^2+^ to sites III/IV in the presence of 1 mM Mg^2+^ resulted in a K_d_ (0.14 ± 0.01 μM) that was not significantly different (p<0.05) than seen for the 3 mM Mg^2+^ preincubation (K_d_ = 0.08 ± 0.01 μM) (**Figure 5 and Table S3**).

At sites III/IV, for the 1 mM Mg^2+^ pre-incubation, the interaction proceeded with favorable enthalpy (ΔH = −6.87 ± 0.09 kcal*mol^-1^) and entropy (T*ΔS = 2.50 ± 0.10 kcal*mol^-1^). For the 3 mM Mg^2+^ condition, the reaction was again exothermic (ΔH = −6.19 ± 0.06 kcal*mol^-1^) with a positive change in T*ΔS (3.50 ± 0.06 kcal*mol^-1^).

## Discussion

This study provides novel information regarding the thermodynamics that underlie the interaction between cTnC and two physiologically prevalent divalent cations. Our results significantly advance the understanding of the mechanisms and role of modifications in cellular Mg^2+^ in control of the cTnC Ca^2+^ switch. Cellular free Mg^2+^ is known to change in pathological conditions in the heart, but mechanisms of its effects on activation of the myofilaments remains incompletely understood.

As seen in previous reports, we found the binding of Ca^2+^ to N-cTnC to be driven by entropy and unfavorable enthalpy (**Table S2**) (Stevens, Rayani et al. 2017, Johnson, Fulcher et al. 2019). The favorable ΔS may be due in part to the hydration enthalpy of Ca^2+^ which is thought to be on the order of ~375 kcal*mol^-1^ and slightly lower than that of Mg^2+^ (~460 kcal*mol^-1^) (Smith DW, 1977). It is also possible that the endothermic nature of these interactions results from other factors such as the exchange of protons that are transferred from the ligand to the buffer upon Ca^2+^ binding (Skowronsky, Schroeter et al. 2013).

Measurement of Ca^2+^ binding to cTnC is often achieved indirectly by measuring the fluorescence change and correlating this to the conformational change that results from the interaction. Fluorescent molecules such as IAANS (Wang, Huang et al. 1997, Hazard, Kohout et al. 1998, Li, Stevens et al. 2013) or reporters such as F27W (Gillis, Blumenschein et al. 2003) can be used to quantify this binding interaction. At 21°C bovine F27W cTnC had a K_d_ of ~5 μM and IAANS labelled C35S cTnC had a K_d_ of ~7 μM (Gillis, Marshall et al. 2000, Tikunova and Davis 2004). Through fluorescence-based measurement, the K_d_ of N-cTnC for Ca^2+^ was previously reported to be between 11.3 μM – 12.3 μM (Liang, Chung et al. 2008, Pinto, Parvatiyar et al. 2009). These parameters agree with our measured Ca^2+^ binding to apo-state N-cTnC and apo-state cTnC (**Table S2**), deviating only slightly due to buffer and temperature conditions.

Normally, cytosolic [Mg^2+^]_free_ is maintained around ~0.5 – 1 mM (Romani 2011). At these concentrations, Mg^2+^ is known to compete with Ca^2+^ for sites III and IV. Circular dichroism has been used to show that Ca^2+^ binding to sites III/IV increases the α-helical content of cTnC, from 19 to 41 percent (Herzberg and James 1985, Yumoto, Nara et al. 2001) and causes conformational changes that remove non-polar amino acids from the solvent exposed environment (Sturtevant 1977). This contrasts with NMR-based visualization of N-cTnC, in which the apo-state and Ca^2+^-bound forms showed minimal structural deviation (Spyracopoulos, Li et al. 1997).

Ca^2+^ has many times greater polarizability than Mg^2+^ and lower hydration energy (Carafoli and Krebs 2016). The bare ion radius of Mg^2+^ (0.65 Å) is smaller than Ca^2+^ (0.99 Å) (Lockless, Zhou et al. 2007), conversely, in its hydrated form Mg^2+^ (4.3 Å) is larger than Ca^2+^ (4.1 Å) (Maguire 2006). In other Ca^2+^ binding proteins such as calmodulin (CaM), metals with similar ionic radii are able to substitute for this cation (Chao, Suzuki et al. 1984, Malmendal, Linse et al. 1999). Mg^2+^ is able to bind to CaM, but does not induce the conformational change associated with Ca^2+^ binding; a phenomenon that is commonly observed in cell biology (Follenius and Gerard 1984, Gilli, Lafitte et al. 1998).

Normally, six oxygen atoms arranged in an octahedral geometry are thought to coordinate Mg^2+^ (Linse and Forsen 1995). This is one less oxygen than needed to coordinate Ca^2+^ through a pentagonal bipyramid (Cates, Berry et al. 1999). However, Ca^2+^ can be coordinated by 6 – 8 coordinating residues (but also by as many as 12) at a distance that can vary greatly (2.3 – 2.7 Å) compared to a much smaller variance for Mg^2+^ coordination (2.0 – 2.2 Å) (Brini, Cali et al. 2012).

Ca^2+^ and Mg^2+^ are most often coordinated by oxygen atoms, this is usually a hydroxyl group for Mg^2+^ and a carboxyl group for Ca^2+^ (Harding 2002). Ca^2+^ is most frequently coordinated by side chains of aspartic acid, glutamic acid, asparagine, followed by serine/threonine, while Mg^2+^ is most frequently coordinated by aspartic acid, glutamic acid, histidine, threonine, serine, and asparagine. (Dokmanic, Sikic et al. 2008). EF hand-containing proteins have also been shown to bind Mg^2+^ when there are appropriately placed negatively charged amino residues (especially in the +z and – z positions) (Reid and Procyshyn 1995, Tikunova, Black et al. 2001, Davis, Rall et al. 2002). In site II of mammalian cTnC, there is a polar serine at the +z position (residue 69) and a negatively charged glutamic acid at the –z position (residue 76) (**Figure S1**).

Data from earlier studies suggested Mg^2+^ bound exclusively at sites III and IV of TnC (Potter and Gergely 1975). Shortly thereafter, a limited series of equilibrium dialysis experiements did not show competition between Mg^2+^ and Ca^2+^ for the N-terminal sites of cTnC, instead, other binding sites were suggested (Holroyde, Robertson et al. 1980). Later still, enthalpic titrations were unable to visualize a discernable change in Mg^2+^ binding to the low affinity sites of skeletal TnC (Yamada and Kometani 1982, Li, Chandra et al. 1994). However, assuming competitive binding fluorescence assays at room temperature determined the K_d_ associated with Mg^2+^ binding to be about 4 mM (Johnson, Collins et al. 1980). More recently, Ca^2+^ sensitivity of the actomyosin ATPase and force production of skinned rat cardiac cells was unaltered when Mg^2+^ was increased from 1 to 8 mM (Allen, Xu et al. 2000). These findings were brought into question, however, by studies that utilized metallochromic indicators to deduce sufficiently high Mg^2+^-affinity at the regulatory sites of skeletal TnC (Ogawa 1985).

The observation of Mg^2+^ binding to the low affinity sites of N-cTnC has led to the suggestion that differences in affinity may be due at least in part to the Ca^2+^ buffering and thus the free concentration of the ion in these experiments. Given the kinetic rates associated with these interactions, it is difficult to have confidence in EGTA determined rates of binding (Ebashi and Ogawa 1988). Moreover, given the temperature sensitivity of cTnC, this factor alone can alter experimental outcomes by orders of magnitude (Kohama 1980, Gillis, Moyes et al. 2003). Change in sensitivity in the face of altered temperature has been suggested to result mostly from binding to the low affinity sites and possibly through interactions with other members of the Tn complex (Godt and Lindley 1982, Stephenson and Williams 1982, Wnuk, Schoechlin et al. 1984).

Experiments testing the effects of alterations in free Mg^2+^ on Ca^2+^-activation of isolated myofibrils and skinned fiber bundles from different laboratories provide corroborative findings supporting the credibility of our postulate of a role for cytosolic Mg^2+^ as a controller of cTnC function at the N-lobe. Fabiato and Fabiato showed that increasing concentration of free Mg^2+^ decreases myofilament Ca^2+^ sensitivity of skinned cardiomyocytes (Fabiato and Fabiato 1975). [Mg^2+^] affects the Ca^2+^-sensitivity of the myofibrillar ATPase as well as actomyosin tension development in both skeletal and cardiac muscle preparations (Donaldson and Kerrick 1975, Kerrick and Donaldson 1975, Solaro and Shiner 1976, Ashley and Moisescu 1977, Best, Donaldson et al. 1977, Donaldson, Best et al. 1978, Ebashi, Nonomura et al. 1980, Morimoto 1991).

Mg^2+^ affinity of sites III/IV alone is not sufficient to fully explain the change in the force-pCa relationship caused by Mg^2+^ in skinned skeletal muscle fibers (Ebashi and Endo 1968). In rabbit fast skeletal muscle, Mg^2+^ competes with Ca^2+^ for low affinity binding sites of TnC where it binds with an affinity of 1.9*10^2^ M^-1^ (much lower than the 6.2*10^6^ M^-1^ seen for Ca^2+^). The K_A_ associated with sites III and IV was measured to be 1.2*10^6^ M^-1^ for Ca^2+^ and 1.1*10^2^ M^-1^ for Mg^2+^ in canine ventricular skinned myocytes (Pan and Solaro 1987).

In isolated cTnC, Mg^2+^ was found to interact with site II of cTnC with an apparent binding constant of 5.2*10^2^ M^-1^. This was only slightly lower than the constant associated with Mg^2+^ binding to sites III/IV (~10^3^ M^-1^), Ca^2+^ binding to sites III/IV (~10^6^ M^-1^), and Ca^2+^ binding to site II (~10^4^ M^-1^) (Ogawa 1985).

Fluorescence was used to measure the Mg^2+^ affinity of site II at 15 °C (~1.2-1.9 mM) (Tikunova and Davis 2004). In the presence of 3 mM Mg^2+^, the K_d_ associated with binding of Ca^2+^ to site II of full length cTnC was increased from 7 μM in the apo-state to 24 μM (Tikunova and Davis 2004). Moreover, a system containing cTnC-cTnI had 2.5-fold lower Ca^2+^ affinity in the presence of 3 mM Mg^2+^ (Siddiqui, Tikunova et al. 2016). Given these affinities, Tikunova and Davis hypothesized that site II would be 33-44% saturated by 1 mM cytosolic Mg^2+^ at diastolic Ca^2+^ concentrations (Tikunova and Davis 2004).

In a recent ITC study, the Mg^2+^ binding affinity of site II in lobster TnC isoforms, which are similar in sequence to human variants, was explored. Mg^2+^ affinity of site II was a single order of magnitude lower than that of Ca^2+^, such that the cations would compete for binding (Tanaka, Takahashi et al. 2013) under physiological conditions.

In our experiments on N-cTnC and full length cTnC, site II binding affinity of Mg^2+^ was an order of magnitude lower than seen for Ca^2+^ (**Figures 2 and 6; Table S2 and S3**). At these affinities and given the relatively high cytosolic [Mg^2+^]_free_ (Linse and Forsen 1995, Malmendal, Linse et al. 1999), this cation would compete for binding to site II of cTnC (Nara, Morii et al. 2013). Competition experiements were also in agreement (**Figures 1 and 2**) as were studies that utilized a double mutant removing coordinating residues in site II (**Figures 3 and 4**).

In order to further validate the ITC data, we also performed thermodynamic integration (TI) to calculate absolute binding affinities computationally. We performed these calculations for both Ca^2+^ and Mg^2+^ binding separately for both WT N-cTnC and D67A/D73A N-cTnC. For both sets of simulations, the structure of Ca^2+^-bound N-cTnC (PDB:1AP4) was used as the starting parameter and restrained throughout the simulation. ITC measures the thermodynamically quantifiable closed-to-open transition of the N-cTnC molecule. TI does not allow for such a transition, rather, it quantifies only the binding interaction. In the future, the closed structure of N-cTnC (PDB:1SPY) can be simulated to quantify the presumably lower affinity it has for each of Ca^2+^ and Mg^2+^. The difference between these sets of simulations could then be used to better corroborate the ITC data.

For Ca^2+^ binding, our TI results agreed very well with the binding affinities from ITC. For Mg^2+^ binding, the calculated absolute binding affinities were consistently underestimated by about 4 kcal*mol^-1^ but showed the same relative trends. Mg^2+^ was calculated to bind weaker than Ca^2+^ and bind weaker for the D67A/D73A mutation similarly to ITC. The Mg^2+^ absolute binding affinities were likely underestimated for multiple reasons. First, the crystal structure of WT N-cTnC (1AP4) contained Ca^2+^ bound and no structure of Mg^2+^ bound WT N-cTnC was available. We attempted to correct for this issue by minimizing the structure with Mg^2+^ bound. Due to the lack of an exact starting structure and restraints chosen, there is still likely some error. Additionally, while we did try to choose the most accurate Mg^2+^ parameters for binding affinity calculation, there are well documented difficulties in free energy calculations for Mg^2+^, most notably that the free energy of solvation (ΔGsolvation) is consistently underestimated (Steinbrecher, Joung et al. 2011, Panteva, Giambasu et al. 2015). Even when using the same Mg^2+^ force field, solvation ΔG values are also known to have large variations for Mg^2+^ depending on the exact simulation parameters used. For example, Panteva et al. and Li et al. both tried to reproduce Mg^2+^ solvation free energy using the same parameters as Åqvist, but saw variations on the order of 20 kcal*mol^-1^ (Ȧqvist 1990). While this may be an extreme example, it illustrates the difficulty in calculation of free energy changes with Mg^2+^ ions involved. Given these potential errors in TI for ΔGsolvation for Mg^2+^, the fact that we still see relatively good agreement with the ITC data for absolute binding affinity of Mg^2+^ helps further validate the *in vitro* results.

The experiments outlined above were designed with the intent to test the hypothesis that: Ca^2+^ and Mg^2+^ both interact with all the functional EF-hand motifs in cTnC. The interaction with sites III and IV has been established for some time (Johnson, Collins et al. 1980), but site II may also bind Mg^2+^. Interestingly, a hypothesis that is reconcilable with our own was initially put forth; that of six binding sites. In this scenario there were: two Ca^2+^ specific sites, two Mg^2+^ specific sites, and two sites that can bind both cations. During these experiments, only the absence of Mg^2+^ allowed for the binding sites in cTnC to be separated into low affinity sites (~10^5^ M^-1^) and high affinity sites (~10^7^ M^-1^) (Potter and Gergely 1975).

Binding of Mg^2+^ to site II is not expected to induce significant structural changes in N-cTnC based on previous Molecular Dynamics simulation data (Spyracopoulos, Li et al. 1997, Skowronsky, Schroeter et al. 2013, Stevens, Rayani et al. 2017). Therefore, it is likely that the favorable ΔS associated with the interaction is due to increased degrees of freedom for the water molecules that would result when stabilizing hydrogen bonds are transferred from the positively charged metal cation and the negatively charged amino acid side chains in the binding site II to the buffered environment (Skowronsky, Schroeter et al. 2013).

Given that the binding of Ca^2+^ to site II of cTnC at systolic Ca^2+^ levels (0.5 - 1.2 μM) strengthens the interaction with cTnI and the rest of the cTn complex and the orders of magnitude difference between binding affinity at varying levels of filament complexity, (Potter and Gergely 1975, Ramos 1999, Pinto, Parvatiyar et al. 2009, Li, Stevens et al. 2013), care must be taken when translating observations at the level of cTnC to more complex systems. Moreover, a further limitation is part of our approach which utilizes the double mutant D67A/D73A. This mutation was able to reduce the binding of both Ca^2+^ (11.9-fold) and Mg^2+^ (1.8-fold) to site II of N-cTnC; however, the impact on binding might be expected to be greater. It is possible that the effect of this double mutant is to reduce the binding of these cations, especially Mg^2+^ through allosteric interactions. In CaM, mutation of Ca^2+^ coordinating residues within the EF-hand can have structural consequences leading to altered binding kinetics (Wang, Brohus et al. 2020); this is conceivable in our double mutant. Similarily, it is possible that the competition observed between Ca^2+^ and Mg^2+^ for binding to site II of cTnC occurs through structural perturbations which follow binding of Mg^2+^ to an allosteric site. Exploration of these limitations in furture studies may shed light on the true nature of these interactions.

Our ITC results strongly suggest that Mg^2+^ binds to site III/IV and also competes with Ca^2+^ for binding to site II. The amount of Mg^2+^ that binds the regulatory site II is likely to be highly dependent on technique, biological system, and buffer conditions. In N-cTnC, occupation of site II by Mg^2+^ was again seen to reduce the amount of Ca^2+^ which was able to bind this protein, at concentrations that may have physiologically relevant consequences under normal conditions and even more so in the face of diseases which alter the Ca^2+^ sensitivity of contraction.

Moreover, increases in cAMP in the cell through α- and β-adrenergic stimulation elicits extrusion of Mg^2+^ from the cell in mammalian tissues (Wolf, Di Francesco et al. 1996, Fagan and Romani 2001, Cefaratti and Romani 2007) including cardiomyocytes (Vormann and Günther 1987, Howarth, Waring et al. 1994). If shown in the heart, both Na^+^-dependent and independent removal of Mg^2+^ from the cytosol under stressful conditions would lower cytosolic presence of this cation. Despite this, free Mg^2+^ does not fluctuate greatly under such stimulation, suggesting that buffered Mg^2+^ is removed from the cell (Amano, Matsubara et al. 2000). Nevertheless, this altered Mg^2+^ pool may affect the subset of ions available to compete with Ca^2+^ for binding to troponin.

Based on our binding experiments and given the previous studies cited herein, Mg^2+^ may also compete with Ca^2+^ in binding to the regulatory site II. Free Ca^2+^ is tightly regulated at rest (~0.1 μM) despite relatively high total cytosolic concentrations (2.1 – 2.6 mM) (Brini, Cali et al. 2012). Mg^2+^ is also abundant in the cell but is less tightly controlled. Binding of both Ca^2+^ and Mg^2+^ to site II is enothermic, and thus driven by entropy. Relative to Ca^2+^, Mg^2+^ binds site II with lower affinity, however at physiological concentrations or elevation of free Mg^2+^, which accompanies states of energy depletion, it may reduce Ca^2+^ binding leading to structural perturbations that modify the contractile function of the myofilament. Conversely, Mg^2+^ can be altered by diseased states such as secondary hyperparathyroidism which results in hypomagnesia and could potentially impact cardiac contractility (Morsy, Dishmon et al. 2017).

## Conclusions

Our studies provide insights regarding the thermodynamics of metal cation binding to cTnC. The interaction of Ca^2+^ and Mg^2+^ with cTnC are characterized by differences consistent with dissimilar ionic radius, number of required coordinating residues, as well as the energic cost of exposing hydrophobicity amino acids to an aqueous environment. In the cell, these differences are functionally necessitated by dissimilar cytosolic prevalence of each cation. Cellular Mg^2+^ is not necessarily prevalent enough to directly regulate contraction and is not thought to cause a conformational change upon binding to cTnC. However, given the affinites we have observed, its occupation of the binding site may restrict Ca^2+^ binding. This competition for binding likely favors Ca^2+^ and is well tolerated, however elevation of free Mg^2+^, which may accompany states of ATP depletion could have relatively significant functional consequence for cardiac force production, for example during ischemic stresses.

## Supplementary Appendix

**Figure S1 -.**
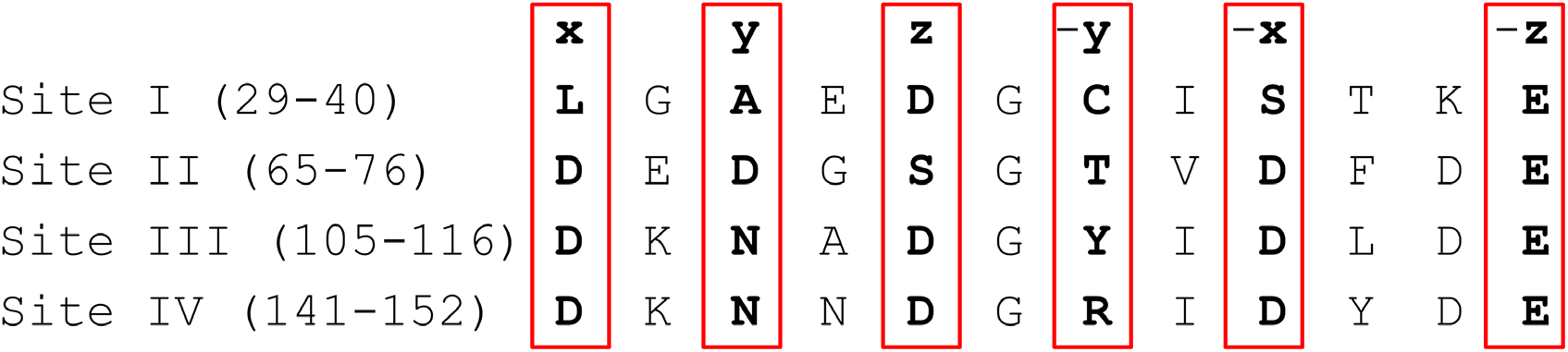
Sequence alignment of the 4 EF hand binding motifs in cTnC. The coordinating residues within EF hands I-IV are shown with the residue number listed in brackets. Each of residues x, y, z, -y, -x, and -z, that make up the helices of the pentagonal bipyramid are indicated. The conservation of the amino acids in each of the coordinating residues between sites II, III, and IV is striking as is the clear differences seen in site I.

**Figure S2 -.**
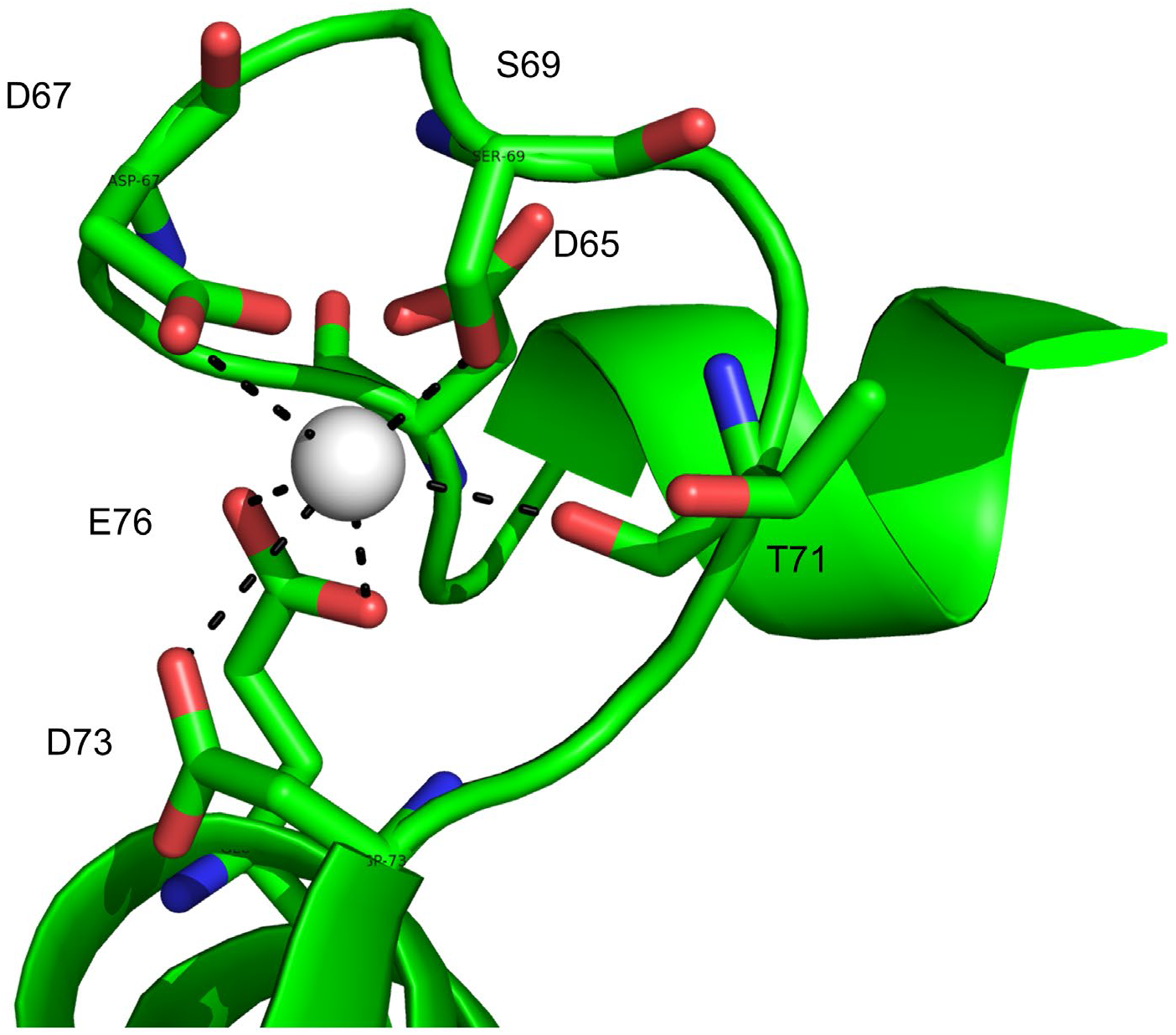
Depiction of site II of WT cTnC coordinating a divalent cation. The D67A/D73A double mutant removes two of the coordinating residues within the EF hand of site II in N-cTnC. The goal of this double mutation was to compare the reduced amount of binding of Ca^2+^ and Mg^2+^ and to gain insight into the locus of binding for each cation. The figure was generated using PyMOL and adapted from the PDB:1J1E x-ray structure.

**Table S1 -.**
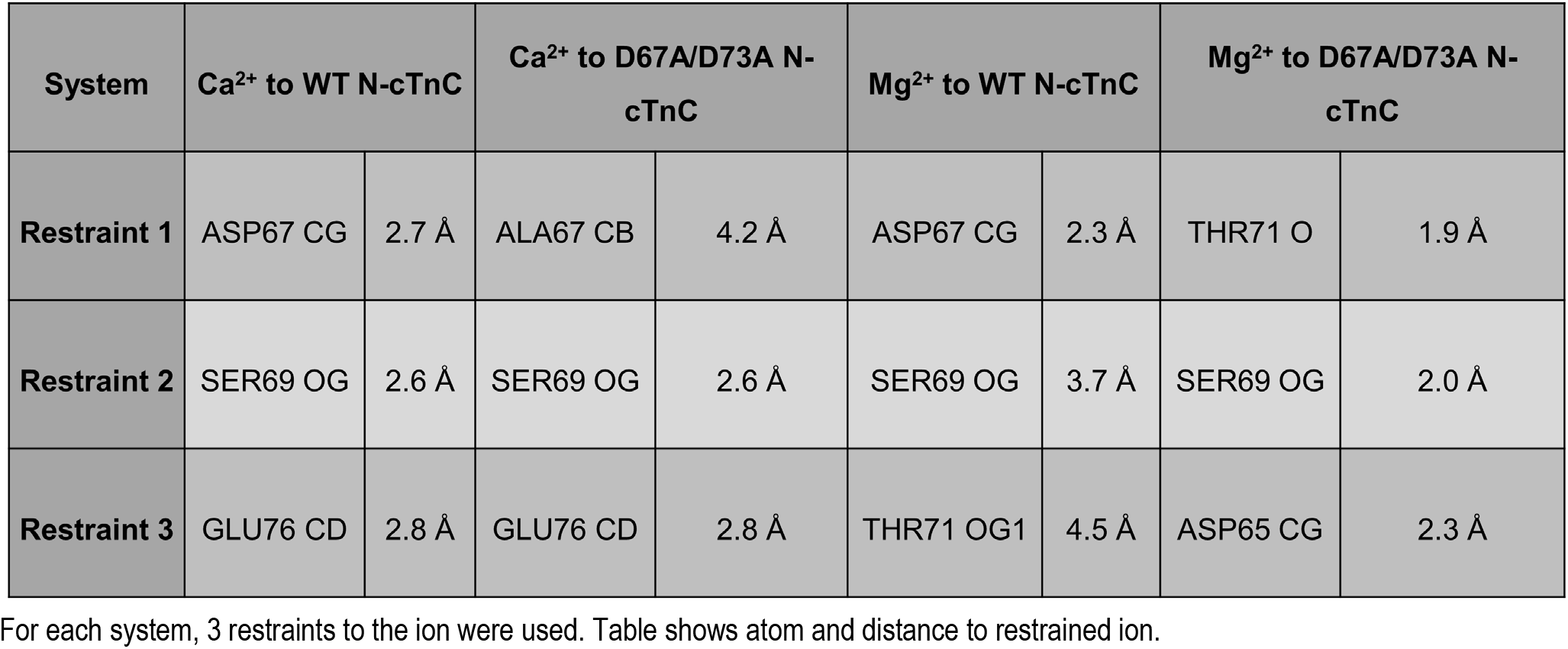
Ion restraints used for thermodynamic integration

| System | Ca<sup>2+</sup> to WT N-cTnC |  | Ca<sup>2+</sup> to D67A/D73A N-cTnC |  | Mg<sup>2+</sup> to WT N-cTnC |  | Mg<sup>2+</sup> to D67A/D73A N-cTnC |  |
| --- | --- | --- | --- | --- | --- | --- | --- | --- |
| **Restraint 1** | ASP67 CG | 2.7 Å | ALA67 CB | 4.2 Å | ASP67 CG | 2.3 Å | THR71 O | 1.9 Å |
| **Restraint 2** | SER69 OG | 2.6 Å | SER69 OG | 2.6 Å | SER69 OG | 3.7 Å | SER69 OG | 2.0 Å |
| **Restraint 3** | GLU76 CD | 2.8 Å | GLU76 CD | 2.8 Å | THR71 OG1 | 4.5 Å | ASP65 CG | 2.3 Å |

For each system, 3 restraints to the ion were used. Table shows atom and distance to restrained ion.

**Table S2 -.** Thermodynamic Parameters for Ca^2+^ and Mg^2+^ binding to N-cTnC

| Titrant | N-cTnC Condition | n | N | K<sub>A</sub>*10<sup>3</sup><br>(M<sup>-1</sup>) | K<sub>d</sub><br>(μM) | ΔH<br>(kcal*mol<sup>-1</sup>) | T*ΔS<br>(kcal*mol<sup>-1</sup>) | ΔG<br>(kcal*mol<sup>-1</sup>) |
| --- | --- | --- | --- | --- | --- | --- | --- | --- |
| Ca<sup>2+</sup> | Apo-State | 7 | 1.01 ± 0.01 | 66.1<sup>A</sup><br>± 2.0 | 15.2<sup>C</sup><br>± 0.5 | 3.82<sup>A</sup><br>± 0.04 | 10.39<sup>A</sup><br>± 0.03 | -6.57<sup>H</sup><br>± 0.02 |
|  | D67A/D73A | 8 | 1.00 | 5.9<sup>E, F</sup><br>± 0.5 | 180.3<sup>C</sup><br>± 16.2 | 0.23<sup>H, I</sup><br>± 0.02 | 5.36<sup>G</sup><br>± 0.05 | -5.12<sup>D</sup><br>± 0.05 |
|  | + 1 mM Mg<sup>2+</sup> | 6 | 1.00 | 31.2<sup>B</sup><br>± 2.7 | 33.1<sup>C</sup><br>± 2.57 | 2.47<sup>B</sup><br>± 0.05 | 8.58<sup>B</sup><br>± 0.01 | -6.12<sup>G</sup><br>± 0.05 |
|  | + 3 mM Mg<sup>2+</sup> | 7 | 1.00 | 20.9<sup>C</sup><br>± 1.3 | 48.9<sup>C</sup><br>± 2.81 | 1.73<sup>C</sup><br>± 0.05 | 7.62<sup>C</sup><br>± 0.03 | -5.89<sup>G</sup><br>± 0.04 |
|  | + 5 mM Mg<sup>2+</sup> | 7 | 1.00 | 14.0<sup>D</sup><br>± 0.7 | 72.3<sup>C</sup><br>± 3.17 | 1.52<sup>D</sup><br>± 0.05 | 7.17<sup>D</sup><br>± 0.04 | -5.65<sup>F</sup><br>± 0.02 |
|  | + 10 mM Mg<sup>2+</sup> | 7 | 1.00 | 9.7<sup>D, E</sup><br>± 0.5 | 105.3<sup>C</sup><br>± 5.54 | 1.13<sup>E</sup><br>± 0.04 | 6.57<sup>E</sup><br>± 0.02 | -5.43<sup>E, F</sup><br>± 0.03 |
|  | + 20 mM Mg<sup>2+</sup> | 5 | 1.00 | 8.2<sup>E</sup><br>± 0.1 | 122.8<sup>C</sup><br>± 2.06 | 0.78<sup>F</sup><br>± 0.04 | 6.13<sup>F</sup><br>± 0.03 | -5.35<sup>D, E</sup><br>± 0.01 |
| Mg<sup>2+</sup> | Apo-State | 5 | 1.00 | 1.54<sup>F, G</sup><br>± 0.06 | 652.8<sup>B, C</sup><br>± 28.4 | 2.64<sup>B</sup><br>± 0.1 | 6.99<sup>D</sup><br>± 0.07 | -4.35<sup>C</sup><br>± 0.03 |
|  | D67A/D73A | 5 | 1.00 | 0.90<sup>F, G</sup><br>± 0.08 | 1148.6<sup>A</sup><br>± 95.0 | 0.76<sup>F</sup><br>± 0.04 | 4.78<sup>H</sup><br>± 0.03 | -4.02<sup>B</sup><br>± 0.05 |
|  | + 1 mM Ca<sup>2+</sup> | 5 | 1.00 | 0.55<sup>G</sup><br>± 0.05 | 1870.0<sup>A</sup><br>± 171.5 | 0.48<sup>G</sup><br>± 0.04 | 4.22<sup>I</sup><br>± 0.03 | -3.73<sup>A</sup><br>± 0.05 |
|  | + 2 mM Ca<sup>2+</sup> | 6 | 1.00 | 0.47<sup>G</sup><br>± 0.05 | 2251.7<sup>A</sup><br>± 261.8 | 0.41<sup>G, H</sup><br>± 0.04 | 4.04<sup>I, J</sup><br>± 0.07 | -3.63<sup>A</sup><br>± 0.07 |
|  | + 3 mM Ca<sup>2+</sup> | 8 | 1.00 | 0.52<sup>G</sup><br>± 0.04 | 2037.5<sup>A</sup><br>± 172.2 | 0.24<sup>H, I</sup><br>± 0.02 | 3.92<sup>J</sup><br>± 0.04 | -3.69<sup>A</sup><br>± 0.05 |
|  | + 5 mM Ca<sup>2+</sup> | 9 | 1.00 | 0.50<sup>G</sup><br>± 0.08 | 2255.8<sup>A</sup><br>± 220.1 | 0.20<sup>I</sup><br>± 0.02 | 3.84<sup>J</sup><br>± 0.06 | -3.64<sup>A</sup><br>± 0.07 |

In this figure, n indicates the experimental repeats and N indicates the stoichiometry number. 4 mM Ca<sup>2+</sup> and 20 mM Mg<sup>2+</sup> titrations into 200 μM N-cTnC were the baseline conditions. For the Ca<sup>2+</sup> titrations, pre-incubation with 1, 3, 5, 10, and 20 mM Mg<sup>2+</sup> prior to titrations was used to study the site II occupation by both cations. The affinity and enthalpy change associated with the interaction was decreased with increasing amounts of Mg<sup>2+</sup> incubated. For the Mg<sup>2+</sup> titrations, pre-incubation with 1, 2, 3, 5 mM Ca<sup>2+</sup> prior to titrations was used to study the binding of both cations to site II. Increasing the concentration of Ca<sup>2+</sup> decreased the binding affinity of Mg<sup>2+</sup> to site II and decreased the enthalpy change, therefore less Mg<sup>2+</sup> binds N-cTnC in the presence of Ca<sup>2+</sup>. The D67A/D73A mutant was used to reduce the binding of both cations to the EF hand of site II, albeit to a different extent for each. All parameters are displayed as mean ± SEM, with the exception of the stoichiometric ratio for the Mg<sup>2+</sup> binding which was constrained to 1.00 to facilitate fitting. Titrations not linked with the same superscripted letter were significantly different (p<0.05). The first letter of the alphabet indicates the largest mean and each subsequent letter denotes a significantly lower mean.

**Table S3 -.** Thermodynamic Parameters for Ca^2+^ and Mg^2+^ binding to full-length cTnC

|  | Titrant | cTnC condition | n | N | K<sub>A</sub><br>*10<sup>3</sup> (M<sup>-1</sup>) | K<sub>d</sub><br>(μM) | ΔH<br>(kcal*mol<sup>-1</sup>) | T*ΔS<br>(kcal*mol<sup>-1</sup>) | ΔG<br>(kcal*mol<sup>-1</sup>) |
| --- | --- | --- | --- | --- | --- | --- | --- | --- | --- |
| Site II | Ca<sup>2+</sup> | Apo-State | 7 | 1.91<br>± 0.07 | 44.3<sup>A</sup><br>± 1.0 | 22.7<sup>B</sup><br>± 0.5 | 3.71<sup>A</sup><br>± 0.06 | 10.0<sup>A</sup><br>± 0.07 | -6.29<sup>C</sup><br>± 0.20 |
|  | Mg<sup>2+</sup> | Apo-State | 8 | 32.9<br>± 0.5 | 2.47<sup>C</sup><br>± 0.05 | 406.1<sup>A</sup><br>± 7.9 | 0.091<sup>D</sup><br>± 0.001 | 4.71<sup>D</sup><br>± 0.01 | -4.62<sup>A</sup><br>± 0.11 |
|  | Ca<sup>2+</sup> | + 1 mM Mg<sup>2+</sup> | 8 | 2.10<br>± 0.05 | 32.4<sup>B</sup><br>± 0.9 | 31.0<sup>B</sup><br>± 0.9 | 1.93<sup>B</sup><br>± 0.09 | 8.09<sup>B</sup><br>± 0.08 | -6.16<sup>B</sup><br>± 0.59 |
|  | Ca<sup>2+</sup> | + 3 mM Mg<sup>2+</sup> | 10 | 2.96<br>± 0.06 | 29.1<sup>B</sup><br>± 1.7 | 35.5<sup>B</sup><br>± 2.1 | 0.75<sup>C</sup><br>± 0.03 | 6.83<sup>C</sup><br>± 0.04 | -6.08<sup>B</sup><br>± 0.54 |
| Sites III/IV | Ca<sup>2+</sup> | Apo-State | 7 | 2.19<br>± 0.05 | 8757.1<sup>B</sup><br>± 951.4 | 0.12<sup>B</sup><br>± 0.02 | -8.20<sup>D</sup><br>± 0.07 | 1.24<sup>D</sup><br>± 0.07 | -9.44<sup>B</sup><br>± 0.61 |
|  | Mg<sup>2+</sup> | Apo-State | 8 | 4.92<br>± 0.15 | 60.5<sup>C</sup><br>± 3.0 | 16.7<sup>A</sup><br>± 0.7 | -0.23<sup>A</sup><br>± 0.01 | 6.28<sup>A</sup><br>± 0.03 | -6.51<sup>A</sup><br>± 0.31 |
|  | Ca<sup>2+</sup> | + 1 mM Mg<sup>2+</sup> | 8 | 2.02<br>± 0.04 | 7526.3<sup>B</sup><br>± 333.5 | 0.14<sup>B</sup><br>± 0.01 | -6.87<sup>C</sup><br>± 0.09 | 2.50<sup>C</sup><br>± 0.10 | -9.37<sup>B</sup><br>± 0.50 |
|  | Ca<sup>2+</sup> | + 3 mM Mg<sup>2+</sup> | 10 | 1.94<br>± 0.04 | 12870.0<sup>A</sup><br>± 939.6 | 0.08<sup>B</sup><br>± 0.01 | -6.19<sup>B</sup><br>± 0.06 | 3.50<sup>B</sup><br>± 0.06 | -9.79<sup>C</sup><br>± 0.26 |

In this figure, n indicates the experimental repeats and N indicates the stoichiometry number. 6 mM Ca<sup>2+</sup> and 40 mM Mg<sup>2+</sup> titrations into 100 μM N-cTnC were the baseline conditions. Ca<sup>2+</sup> was also titrated into protein pre-incubated with 1 and 3 mM Mg<sup>2+</sup> to study the competition for all three binding sites. Binding to site II was analyzed separately from binding to sites III/IV where analysis of variance was used to test for a significant difference in the means for each thermodynamic parameter. At the level of each parameter, Tukey's post hoc test was carried out. The results of this test are indicated by superscripted letters, where conditions with unique letters are significantly different (p < 0.05).

## Notes

### Competing Interest Statement

The authors have declared no competing interest.

## References

Allen, K., Y. Y. Xu and W. G. Kerrick (2000). “Ca2+measurements in skinned cardiac fibers: effects of Mg2+ on Ca2+ activation of force and fiber ATPase.” J Appl Physiol (1985) 88(1): 180–185.

Allen, T. S., L. D. Yates and A. M. Gordon (1992). “Ca2+-dependence of structural changes in troponin-C in demembranated fibers of rabbit psoas muscle.” Biophys J 61(2): 399–409.

Amano, T., T. Matsubara, J. Watanabe, S. Nakayama and N. Hotta (2000). “Insulin modulation of intracellular free magnesium in heart: involvement of protein kinase C.” British journal of pharmacology 130(4): 731–738.

Ȧqvist, J. (1990). “Ion-water interaction potentials derived from free energy perturbation simulations.” The Journal of Physical Chemistry 94(21): 8021–8024.

Ashley, C. C. and D. G. Moisescu (1977). “Effect of changing the composition of the bathing solutions upon the isometric tension-pCa relationship in bundles of crustacean myofibrils.” J Physiol 270(3): 627–652.

Bers, D. M. (2000). “Calcium Fluxes Involved in Control of Cardiac Myocyte Contraction.” Circulation Research 87(4): 275–281.

Best, P. M., S. K. Donaldson and W. G. Kerrick (1977). “Tension in mechanically disrupted mammalian cardiac cells: effects of magnesium adenosine triphosphate.” J Physiol 265(1): 1–17.

Boresch, S., F. Tettinger, M. Leitgeb and M. Karplus (2003). “Absolute Binding Free Energies: A Quantitative Approach for Their Calculation.” The Journal of Physical Chemistry B 107(35): 9535–9551.

Bowman, J. D. and S. Lindert (2018). “Molecular Dynamics and Umbrella Sampling Simulations Elucidate Differences in Troponin C Isoform and Mutant Hydrophobic Patch Exposure.” 122(32): 7874–7883.

Brini, M., T. Cali, D. Ottolini and E. Carafoli (2012). “Calcium pumps: why so many?” Compr Physiol 2(2): 1045–1060.

Carafoli, E. and J. Krebs (2016). “Why Calcium? How Calcium Became the Best Communicator.” J Biol Chem 291(40): 20849–20857.

Cates, M. S., M. B. Berry, E. L. Ho, Q. Li, J. D. Potter and G. N. Phillips, Jr. (1999). “Metal-ion affinity and specificity in EF-hand proteins: coordination geometry and domain plasticity in parvalbumin.” Structure 7(10): 1269–1278.

Cefaratti, C. and A. M. Romani (2007). “Functional characterization of two distinct Mg 2+ extrusion mechanisms in cardiac sarcolemmal vesicles.” Molecular and cellular biochemistry 303(1-2): 63–72.

Chao, S. H., Y. Suzuki, J. R. Zysk and W. Y. Cheung (1984). “Activation of calmodulin by various metal cations as a function of ionic radius.” Mol Pharmacol 26(1): 75–82.

Cheung, J. Y., D. L. Tillotson, R. Yelamarty and R. Scaduto (1989). “Cytosolic free calcium concentration in individual cardiac myocytes in primary culture.” American Journal of Physiology-Cell Physiology 256(6): C1120–C1130.

Dai, L. J., P. A. Friedman and G. A. Quamme (1997). “Cellular mechanisms of chlorothiazide and cellular potassium depletion on Mg2+ uptake in mouse distal convoluted tubule cells.” Kidney Int 51(4): 1008–1017.

Davis, J. P., J. A. Rall, P. J. Reiser, L. B. Smillie and S. B. Tikunova (2002). “Engineering competitive magnesium binding into the first EF-hand of skeletal troponin C.” J Biol Chem 277(51): 49716–49726.

Dokmanic, I., M. Sikic and S. Tomic (2008). “Metals in proteins: correlation between the metal-ion type, coordination number and the amino-acid residues involved in the coordination.” Acta Crystallogr D Biol Crystallogr 64(Pt 3: 257–263.

Donaldson, S. K., P. M. Best and G. L. Kerrick (1978). “Characterization of the effects of Mg2+ on Ca2+- and Sr2+-activated tension generation of skinned rat cardiac fibers.” J Gen Physiol 71(6): 645–655.

Donaldson, S. K. and W. G. Kerrick (1975). “Characterization of the effects of Mg2+ on Ca2+- and Sr2+-activated tension generation of skinned skeletal muscle fibers.” J Gen Physiol 66(4): 427–444.

Ebashi, S. and M. Endo (1968). “Calcium ion and muscle contraction.” Prog Biophys Mol Biol 18: 123–183.

Ebashi, S., Y. Nonomura, K. Kohama, T. Kitazawa and T. Mikawa (1980). “Regulation of muscle contraction by Ca ion.” Mol Biol Biochem Biophys 32: 183–194.

Ebashi, S. and Y. Ogawa (1988). “Ca2+ in contractile processes.” Biophys Chem 29(1-2): 137–143.

Fabiato, A. and F. Fabiato (1975). “Effects of magnesium on contractile activation of skinned cardiac cells.” J Physiol 249(3): 497–517.

Fagan, T. E. and A. Romani (2001). “α1-Adrenoceptor-induced Mg2+ extrusion from rat hepatocytes occurs via Na+-dependent transport mechanism.” American Journal of Physiology-Gastrointestinal and Liver Physiology 280(6): G1145–G1156.

Farah, C. S. and F. C. Reinach (1995). “The troponin complex and regulation of muscle contraction.” Faseb j 9(9): 755–767.

Filatov, V. L., A. G. Katrukha, T. V. Bulargina and N. B. Gusev (1999). “Troponin: structure, properties, and mechanism of functioning.” Biochemistry (Mosc) 64(9): 969–985.

Follenius, A. and D. Gerard (1984). “Fluorescence investigations of calmodulin hydrophobic sites.” Biochem Biophys Res Commun 119(3): 1154–1160.

Francois, J. M., C. Gerday, F. G. Prendergast and J. D. Potter (1993). “Determination of the Ca2+ and Mg2+ affinity constants of troponin C from eel skeletal muscle and positioning of the single tryptophan in the primary structure.” J Muscle Res Cell Motil 14(6): 585–593.

Gifford, Jessica L., Michael P. Walsh and Hans J. Vogel (2007). “Structures and metal-ion-binding properties of the Ca<sup>2+</sup>-binding helix–loop–helix EF-hand motifs.” Biochemical Journal 405(2): 199–221.

Gilli, R., D. Lafitte, C. Lopez, M. Kilhoffer, A. Makarov, C. Briand and J. Haiech (1998). “Thermodynamic analysis of calcium and magnesium binding to calmodulin.” Biochemistry 37(16): 5450–5456.

Gillis, T. E., T. M. Blumenschein, B. D. Sykes and G. F. Tibbits (2003). “Effect of temperature and the F27W mutation on the Ca2+ activated structural transition of trout cardiac troponin C.” Biochemistry 42(21): 6418–6426.

Gillis, T. E., C. R. Marshall, X.-H. Xue, T. J. Borgford and G. F. Tibbits (2000). “Ca2+ binding to cardiac troponin C: effects of temperature and pH on mammalian and salmonid isoforms.” American Journal of Physiology-Regulatory, Integrative and Comparative Physiology 279(5): R1707–R1715.

Gillis, T. E., C. D. Moyes and G. F. Tibbits (2003). “Sequence mutations in teleost cardiac troponin C that are permissive of high Ca 2+ affinity of site II.” American Journal of Physiology-Cell Physiology 284(5): C1176–C1184.

Godt, R. E. (1974). “Calcium-activated tension of skinned muscle fibers of the frog. Dependence on magnesium adenosine triphosphate concentration.” J Gen Physiol 63(6): 722–739.

Godt, R. E. and B. D. Lindley (1982). “Influence of temperature upon contractile activation and isometric force production in mechanically skinned muscle fibers of the frog.” J Gen Physiol 80(2): 279–297.

Godt, R. E. and J. L. Morgan (1984). “Contractile responses to MgATP and pH in a thick filament regulated muscle: studies with skinned scallop fibers.” Adv Exp Med Biol 170: 569–572.

Grossoehme, N. E., A. M. Spuches and D. E. Wilcox (2010). “Application of isothermal titration calorimetry in bioinorganic chemistry.” J Biol Inorg Chem 15(8): 1183–1191.

Harding, M. M. (2002). “Metal-ligand geometry relevant to proteins and in proteins: sodium and potassium.” Acta Crystallogr D Biol Crystallogr 58(Pt 5): 872–874.

Hazard, A. L., S. C. Kohout, N. L. Stricker, J. A. Putkey and J. J. Falke (1998). “The kinetic cycle of cardiac troponin C: calcium binding and dissociation at site II trigger slow conformational rearrangements.” Protein Science 7(11): 2451–2459.

Herzberg, O. and M. N. James (1985). “Structure of the calcium regulatory muscle protein troponin-C at 2.8 Å resolution.”

Holroyde, M., S. Robertson, J. Johnson, R. Solaro and J. Potter (1980). “The calcium and magnesium binding sites on cardiac troponin and their role in the regulation of myofibrillar adenosine triphosphatase.” Journal of Biological Chemistry 255(24): 11688–11693.

Hongo, K., M. Konishi and S. Kurihara (1994). “Cytoplasmic free Mg2+ in rat ventricular myocytes studied with the fluorescent indicator furaptra.” Jpn J Physiol 44(4): 357–378.

Houdusse, A., M. L. Love, R. Dominguez, Z. Grabarek and C. Cohen (1997). “Structures of four Ca2+-bound troponin C at 2.0 A resolution: further insights into the Ca2+-switch in the calmodulin superfamily.” Structure 5(12): 1695–1711.

Howarth, F., J. Waring, B. Hustler and J. Singh (1994). “Effects of extracellular magnesium and beta adrenergic stimulation on contractile force and magnesium mobilization in the isolated rat heart.” Magnesium research 7(3-4): 187–197.

Johnson, J. D., S. C. Charlton and J. D. Potter (1979). “A fluorescence stopped flow analysis of Ca2+ exchange with troponin C.” J Biol Chem 254(9): 3497–3502.

Johnson, J. D., J. H. Collins, S. P. Robertson and J. D. Potter (1980). “A fluorescent probe study of Ca2+ binding to the Ca2+-specific sites of cardiac troponin and troponin C.” Journal of Biological Chemistry 255(20): 9635–9640.

Johnson, J. D., R. J. Nakkula, C. Vasulka and L. B. Smillie (1994). “Modulation of Ca2+ exchange with the Ca(2+)-specific regulatory sites of troponin C.” J Biol Chem 269(12): 8919–8923.

Johnson, R. A., L. M. Fulcher, K. Vang, C. D. Palmer, N. E. Grossoehme and A. M. Spuches (2019). “In depth, thermodynamic analysis of Ca(2+) binding to human cardiac troponin C: Extracting buffer-independent binding parameters.” Biochim Biophys Acta Proteins Proteom 1867(4): 359–366.

Kawasaki, Y. and J.-P. van Eerd (1972). “The effect of Mg++ on the conformation of the Ca++-binding component of troponin.” Biochemical and Biophysical Research Communications 49(4): 898–905.

Kerrick, W. G. L. and S. K. B. Donaldson (1975). “The comparative effects of [Ca2+] and [Mg2+] on tension generation in the fibers of skinned frog skeletal muscle and mechanically disrupted rat ventricular cardiac muscle.” Pflügers Archiv 358(3): 195–201.

Kirschenlohr, H. L., A. A. Grace, J. I. Vandenberg, J. C. Metcalfe and G. A. Smith (2000). “Estimation of systolic and diastolic free intracellular Ca2+ by titration of Ca2+ buffering in the ferret heart.” Biochem J 346 Pt 2: 385–391.

Kohama, K. (1980). “Role of the high affinity Ca binding sites of cardiac and fast skeletal troponins.” J Biochem 88(2): 591–599.

Kometani, K. and K. Yamada (1983). “Enthalpy, entropy and heat capacity changes induced by binding of calcium ions to cardiac troponin C.” Biochemical and biophysical research communications 114(1): 162–167.

L DeLano, W. (2002). The PyMOL Molecular Graphics System (2002) DeLano Scientific, Palo Alto, CA, USA. http://www.pymol.org.

Laires, M. J., C. P. Monteiro and M. Bicho (2004). “Role of cellular magnesium in health and human disease.” Front Biosci 9: 262–276.

Leavis, P. and E. L. Kraft (1978). “Calcium binding to cardiac troponin C.” Archives of biochemistry and biophysics 186(2): 411–415.

Leelananda, S. P. and S. Lindert (2016). “Computational methods in drug discovery.” Beilstein J Org Chem 12: 2694–2718.

Lewit-Bentley, A. and S. Rety (2000). “EF-hand calcium-binding proteins.” Curr Opin Struct Biol 10(6): 637–643.

Li, A. Y., C. M. Stevens, B. Liang, K. Rayani, S. Little, J. Davis and G. F. Tibbits (2013). “Familial hypertrophic cardiomyopathy related cardiac troponin C L29Q mutation alters length-dependent activation and functional effects of phosphomimetic troponin I*.”

Li, M. X., M. Chandra, J. R. Pearlstone, K. I. Racher, G. Trigo-Gonzalez, T. Borgford, C. M. Kay and L. B. Smillie (1994). “Properties of isolated recombinant N and C domains of chicken troponin C.” Biochemistry 33(4): 917–925.

Li, M. X., L. Spyracopoulos and B. D. Sykes (1999). “Binding of Cardiac Troponin-I147-163 Induces a Structural Opening in Human Cardiac Troponin-C.” Biochemistry 38(26): 8289–8298.

Li, P., B. P. Roberts, D. K. Chakravorty and K. M. Merz, Jr. (2013). “Rational Design of Particle Mesh Ewald Compatible Lennard-Jones Parameters for +2 Metal Cations in Explicit Solvent.” J Chem Theory Comput 9(6): 2733–2748.

Liang, B., F. Chung, Y. Qu, D. Pavlov, T. E. Gillis, S. B. Tikunova, J. P. Davis and G. F. Tibbits (2008). “Familial hypertrophic cardiomyopathy-related cardiac troponin C mutation L29Q affects Ca2+ binding and myofilament contractility.” Physiological genomics 33(2): 257–266.

Linse, S. and S. Forsen (1995). “Determinants that govern high-affinity calcium binding.” Adv Second Messenger Phosphoprotein Res 30: 89–151.

Lockless, S. W., M. Zhou and R. MacKinnon (2007). “Structural and thermodynamic properties of selective ion binding in a K+ channel.” PLoS Biol 5(5): e121.

Maguire, M. E. (2006). “Magnesium transporters: properties, regulation and structure.” Front Biosci 11: 3149–3163.

Maier, J. A., C. Martinez, K. Kasavajhala, L. Wickstrom, K. E. Hauser and C. Simmerling (2015). “ff14SB: Improving the Accuracy of Protein Side Chain and Backbone Parameters from ff99SB.” J Chem Theory Comput 11(8): 3696–3713.

Malmendal, A., S. Linse, J. Evenas, S. Forsen and T. Drakenberg (1999). “Battle for the EF-hands: magnesium-calcium interference in calmodulin.” Biochemistry 38(36): 11844–11850.

Morimoto, S. (1991). “Effect of myosin cross-bridge interaction with actin on the Ca2+-binding properties of troponin C in fast skeletal myofibrils.” The Journal of Biochemistry 109(1): 120–126.

Morsy, M. S., D. A. Dishmon, N. Garg and K. T. Weber (2017). “Secondary hyperparathyroidism in heart failure.” The American journal of the medical sciences 354(4): 335–338.

Murray, A. C. and C. M. Kay (1972). “Hydrodynamic and optical properties of troponin A. Demonstration of a conformational change upon binding calcium ion.” Biochemistry 11(14): 2622–2627.

Nara, M., H. Morii and M. Tanokura (2013). “Infrared study of synthetic peptide analogues of the calcium-binding site III of troponin C: The role of helix F of an EF-hand motif.” Biopolymers 99(5): 342–347.

Ogawa, Y. (1985). “Calcium binding to troponin C and troponin: effects of Mg2+, ionic strength and pH.” The Journal of Biochemistry 97(4): 1011–1023.

Pan, B. S. and R. J. Solaro (1987). “Calcium-binding properties of troponin C in detergent-skinned heart muscle fibers.” J Biol Chem 262(16): 7839–7849.

Panteva, M. T., G. M. Giambasu and D. M. York (2015). “Comparison of structural, thermodynamic, kinetic and mass transport properties of Mg(2+) ion models commonly used in biomolecular simulations.” J Comput Chem 36(13): 970–982.

Parmacek, M. S. and R. J. Solaro (2004). “Biology of the troponin complex in cardiac myocytes.” Prog Cardiovasc Dis 47(3): 159–176.

Pinto, J. R., M. S. Parvatiyar, M. A. Jones, J. Liang, M. J. Ackerman and J. D. Potter (2009). “A functional and structural study of troponin C mutations related to hypertrophic cardiomyopathy.” J Biol Chem 284(28): 19090–19100.

Potter, J. D. and J. Gergely (1975). “The calcium and magnesium binding sites on troponin and their role in the regulation of myofibrillar adenosine triphosphatase.” Journal of Biological Chemistry 250(12): 4628–4633.

Potter, J. D., S. P. Robertson and J. D. Johnson (1981). “Magnesium and the regulation of muscle contraction.” Fed Proc 40(12): 2653–2656.

Ramos, C. H. (1999). “Mapping subdomains in the C-terminal region of troponin I involved in its binding to troponin C and to thin filament.” J Biol Chem 274(26): 18189–18195.

Reid, R. E. and R. M. Procyshyn (1995). “Engineering magnesium selectivity in the helix-loop-helix calcium-binding motif.” Arch Biochem Biophys 323(1): 115–119.

Robertson, S., J. D. Johnson and J. Potter (1981). “The time-course of Ca2+ exchange with calmodulin, troponin, parvalbumin, and myosin in response to transient increases in Ca2+.” Biophysical journal 34(3): 559.

Rocklin, G. J., D. L. Mobley, K. A. Dill and P. H. Hunenberger (2013). “Calculating the binding free energies of charged species based on explicit-solvent simulations employing lattice-sum methods: an accurate correction scheme for electrostatic finite-size effects.” J Chem Phys 139(18): 184103.

Romani, A. and A. Scarpa (1992). “Regulation of cell magnesium.” Arch Biochem Biophys 298(1): 1–12.

Romani, A. M. P. (2011). Intracellular magnesium homeostasis. Magnesium in the Central Nervous System. R. Vink and M. Nechifor. Adelaide (AU), University of Adelaide Press (c) 2011 The Authors.

Sacco, C., R. A. Skowronsky, S. Gade, J. M. Kenney and A. M. Spuches (2012). “Calorimetric investigation of copper(II) binding to Abeta peptides: thermodynamics of coordination plasticity.” J Biol Inorg Chem 17(4): 531–541.

Schober, T., S. Huke, R. Venkataraman, O. Gryshchenko, D. Kryshtal, H. S. Hwang, F. J. Baudenbacher and B. C. Knollmann (2012). “Myofilament Ca sensitization increases cytosolic Ca binding affinity, alters intracellular Ca homeostasis, and causes pause-dependent Ca-triggered arrhythmia.” Circulation research 111(2): 170–179.

Seamon, K. B., D. J. Hartshorne and A. A. Bothner-By (1977). “Ca2+ and Mg2+ dependent conformations of troponin C as determined by 1H and 19F nuclear magnetic resonance.” Biochemistry 16(18): 4039–4046.

She, M., W. J. Dong, P. K. Umeda and H. C. Cheung (1998). “Tryptophan mutants of troponin C from skeletal muscle: an optical probe of the regulatory domain.” European journal of biochemistry 252(3): 600–607.

Sia, S. K., M. X. Li, L. Spyracopoulos, S. M. Gagné, W. Liu, J. A. Putkey and B. D. Sykes (1997). “Structure of cardiac muscle troponin C unexpectedly reveals a closed regulatory domain.” Journal of Biological Chemistry 272(29): 18216–18221.

Siddiqui, J. K., S. B. Tikunova, S. D. Walton, B. Liu, M. Meyer, P. P. de Tombe, N. Neilson, P. M. Kekenes-Huskey, H. E. Salhi, P. M. Janssen, B. J. Biesiadecki and J. P. Davis (2016). “Myofilament Calcium Sensitivity: Consequences of the Effective Concentration of Troponin I.” Front Physiol 7: 632.

Skowronsky, R. A., M. Schroeter, T. Baxley, Y. Li, J. M. Chalovich and A. M. Spuches (2013). “Thermodynamics and molecular dynamics simulations of calcium binding to the regulatory site of human cardiac troponin C: evidence for communication with the structural calcium binding sites.” JBIC Journal of Biological Inorganic Chemistry 18(1): 49–58.

Slupsky, C. M. and B. D. Sykes (1995). “NMR solution structure of calcium-saturated skeletal muscle troponin C.” Biochemistry 34(49): 15953–15964.

Solaro, R. J. and J. S. Shiner (1976). “Modulation of Ca2+ control of dog and rabbit cardiac myofibrils by Mg2+. Comparison with rabbit skeletal myofibrils.” Circ Res 39(1): 8–14.

Spyracopoulos, L., M. X. Li, S. K. Sia, S. M. Gagné, M. Chandra, R. J. Solaro and B. D. Sykes (1997). “Calcium-induced structural transition in the regulatory domain of human cardiac troponin C.” Biochemistry 36(40): 12138–12146.

Steinbrecher, T., I. Joung and D. A. Case (2011). “Soft-core potentials in thermodynamic integration: comparing one- and two-step transformations.” J Comput Chem 32(15): 3253–3263.

Stephenson, D. G. and D. A. Williams (1982). “Effects of sarcomere length on the force-pCa relation in fast- and slow-twitch skinned muscle fibres from the rat.” J Physiol 333: 637–653.

Stevens, C. M., K. Rayani, C. E. Genge, G. Singh, B. Liang, J. M. Roller, C. Li, A. Y. Li, D. P. Tieleman and F. van Petegem (2016). “Characterization of Zebrafish Cardiac and Slow Skeletal Troponin C Paralogs by MD Simulation and ITC.” Biophysical Journal 111(1): 38–49.

Stevens, C. M., K. Rayani, G. Singh, B. Lotfalisalmasi, D. P. Tieleman and G. F. Tibbits (2017). “Changes in the dynamics of the cardiac troponin C molecule explain the effects of Ca2+-sensitizing mutations.” Journal of Biological Chemistry 292(28): 11915–11926.

Strynadka, N. C. and M. N. James (1989). “Crystal structures of the helix-loop-helix calcium-binding proteins.” Annu Rev Biochem 58: 951–998.

Sturtevant, J. M. (1977). “Heat capacity and entropy changes in processes involving proteins.” Proc Natl Acad Sci U S A 74(6): 2236–2240.

Sundaralingam, M., R. Bergstrom, G. Strasburg, S. T. Rao, P. Roychowdhury, M. Greaser and B. C. Wang (1985). “Molecular structure of troponin C from chicken skeletal muscle at 3-angstrom resolution.” Science 227(4689): 945–948.

Tanaka, H., H. Takahashi and T. Ojima (2013). “Ca(2)+-binding properties and regulatory roles of lobster troponin C sites II and IV.” FEBS Lett 587(16): 2612–2616.

Tessman, P. A. and A. Romani (1998). “Acute effect of EtOH on Mg2+ homeostasis in liver cells: evidence for the activation of an Na+/Mg2+ exchanger.” Am J Physiol 275(5): G1106–1116.

Tikunova, S. B., D. J. Black, J. D. Johnson and J. P. Davis (2001). “Modifying Mg2+ binding and exchange with the N-terminal of calmodulin.” Biochemistry 40(11): 3348–3353.

Tikunova, S. B. and J. P. Davis (2004). “Designing calcium-sensitizing mutations in the regulatory domain of cardiac troponin C.” Journal of Biological Chemistry 279(34): 35341–35352.

van Eerd, J. P. and K. Takahshi (1976). “Determination of the complete amino acid sequence of bovine cardiac troponin C.” Biochemistry 15(5): 1171–1180.

Vormann, J. and T. Günther (1987). “Amiloride-sensitive net Mg2+ efflux from isolated perfused rat hearts.” Magnesium 6(4): 220–224.

Wang, K., M. Brohus, C. Holt, M. T. Overgaard, R. Wimmer and F. Van Petegem (2020). “Arrhythmia mutations in calmodulin can disrupt cooperativity of Ca2+ binding and cause misfolding.” The Journal of Physiology 598(6): 1169–1186.

Wang, S.-Q., Y.-H. Huang, K.-S. Liu and Z.-Q. Zhou (1997). “Dependence of myocardial hypothermia tolerance on sources of activator calcium.” Cryobiology 35(3): 193–200.

Wilcox, D. E. (2008). “Isothermal titration calorimetry of metal ions binding to proteins: An overview of recent studies.” Inorganica Chimica Acta 361(4): 857–867.

Wnuk, W., M. Schoechlin and E. A. Stein (1984). “Regulation of actomyosin ATPase by a single calcium-binding site on troponin C from crayfish.” J Biol Chem 259(14): 9017–9023.

Wolf, F. I., A. Di Francesco, V. Covacci, D. Corda and A. Cittadini (1996). “Regulation of intracellular magnesium in ascites cells: Involvement of different regulatory pathways.” Archives of biochemistry and biophysics 331(2): 194–200.

Yamada, K. (1978). “The enthalpy titration of troponin C with calcium.” Biochimica et Biophysica Acta (BBA)-Protein Structure 535(2): 342–347.

Yamada, K. (2003). “Calcium binding to troponin C as a primary step of the regulation of contraction. A microcalorimetric approach.” Adv Exp Med Biol 538: 203–212; discussion 213.

Yamada, K. and K. Kometani (1982). “The changes in heat capacity and entropy of troponin C induced by calcium binding.” The Journal of Biochemistry 92(5): 1505–1517.

Yap, K. L., J. B. Ames, M. B. Swindells and M. Ikura (1999). “Diversity of conformational states and changes within the EF-hand protein superfamily.” Proteins 37(3): 499–507.

Yumoto, F., M. Nara, H. Kagi, W. Iwasaki, T. Ojima, K. Nishita, K. Nagata and M. Tanokura (2001). “Coordination structures of Ca2+ and Mg2+ in Akazara scallop troponin C in solution. FTIR spectroscopy of side-chain COO-groups.” Eur J Biochem 268(23): 6284–6290.

Zot, A. S. and J. D. Potter (1987). “Structural aspects of troponin-tropomyosin regulation of skeletal muscle contraction.” Annual review of biophysics and biophysical chemistry 16(1): 535–559.

